# Accelerated design of *Escherichia coli* reduced genomes using a whole-cell model and machine learning

**DOI:** 10.1101/2023.10.30.564402

**Authors:** Ioana M. Gherman, Kieren Sharma, Joshua Rees-Garbutt, Wei Pang, Zahraa S. Abdallah, Thomas E. Gorochowski, Claire S. Grierson, Lucia Marucci

## Abstract

Whole-cell models (WCMs) are multi-scale computational models that aim to simulate the function of all genes and processes within a cell. This approach is promising for designing genomes tailored for specific tasks. However, a limitation of WCMs is their long runtime. Here, we show how machine learning (ML) surrogates can be used to address this limitation by training them on WCM data to accurately predict cell division. Our ML surrogate achieves a 95% reduction in computational time compared to the original WCM. We then show that the surrogate and a genome-design algorithm can generate an *in silico* reduced *E. coli* cell, where 40% of the genes included in the WCM were removed. The reduced genome is validated using the WCM and interpreted biologically using gene ontology analysis. This approach illustrates how the holistic understanding gained from a WCM can be leveraged for synthetic biology tasks while reducing its runtime.

## Introduction

Whole-cell models (WCMs) are comprehensive computational models that aim to capture the dynamics of all core biomolecular processes and components of a cell, as well as their interactions with the wider environment. The development of whole-cell modeling is considered a “grand challenge of the 21^*st*^ century” and is seen as particularly useful for understanding the complex relationship between the genotype and phenotype of a cell.^1^ The derivation of WCMs has been made possible by the rapidly increasing availability of experimental data and high-throughput assays that allow researchers to gain a more holistic understanding of biological processes. There are currently three published WCMs: one for *Mycoplasma genitalium*,^2^ one for *Escherichia coli*^3, 4^ and one for the minimal synthetic cell JCVI-syn3A.^5^ The *E. coli* WCM is represented mechanistically at both the single-cell^3, 6^ and at the population level.^4, 7^

Numerous biological discoveries have arisen from the development and use of WCMs. For example, some studies used a WCM in conjunction with experimental data to examine the growth rates of *M. genitalium* and make predictions concerning kinetic parameters of specific enzymes.^8^ By cross-validating numerous *E. coli* datasets during the construction of the WCM, scientists identified several inconsistencies in the literature, which when resolved, enabled the model to accurately predict *in vitro* protein half-lives.^3^ Additionally, the *E. coli* WCM has recently been applied to better understand the differences between *in vivo* and *in vitro* measurements of tRNA aminoacylation,^9^ and the ways operons benefit bacteria.^10^ The population-level *E. coli* WCM, which considers many independent cells within a shared environment, has been instrumental in assessing antibiotic responses and identifying subgenerationally expressed genes that impact cell survival.^7^ Design-build-test-learn cycles can also be optimized using WCMs, with the *M. genitalium* WCM being used to predict minimal genomes *in silico*.^11–13^ Furthermore, the relationship between gene knock outs, cellular subsystems and cellular phenotypes has been better understood through a combination of WCM simulations and machine learning-based classifiers.^9, 14^

The comprehensive nature of WCMs enables a deeper understanding of life at the cellular level, but it also makes these models inherently complex. The complexity of WCMs predominantly stems from their need to simulate the integrated function of every gene and molecule in a cell. For example, the *M. genitalium* WCM captures the function of 401 genes and 28 cellular processes that were each modeled independently on a short time-scale and then integrated over longer time-scales.^2^ The more recently published *E. coli* single-cell WCM initially included the functionality of 1214 genes (43% of the organism’s well-annotated genes) and 10 cellular processes (e.g. transcription, translation, RNA/protein degradation, complexation, metabolism).^3^ This model is under continuous development and when this study began the number of genes modeled had risen to 1219. To build the *E. coli* single-cell WCM, the authors of the model identified and curated over 19,119 parameters from the literature and integrated them in a computational model based on over 10,000 interdependent mathematical equations representing different biological systems.^3^ The output of this model consists of approximately 219,000 time series representing the state of every molecule modeled by the WCM over time, the flux of every metabolic reaction, as well as other metrics summarizing the state of the cell (e.g. the proportion of RNA per gene degraded at each time step, total macromolecular masses, the probability of each gene being expressed at every time step, the number of ribosomes bound to each gene). An example subset of the output of the *E. coli* WCM can be seen in the top panel in Figure 1, where we illustrate how the number of cells simulated grows exponentially as cells divide. Notably, in this figure we only illustrate three time series for each cell, whereas the WCM actually outputs approximately 219,000.

**Figure 1:**
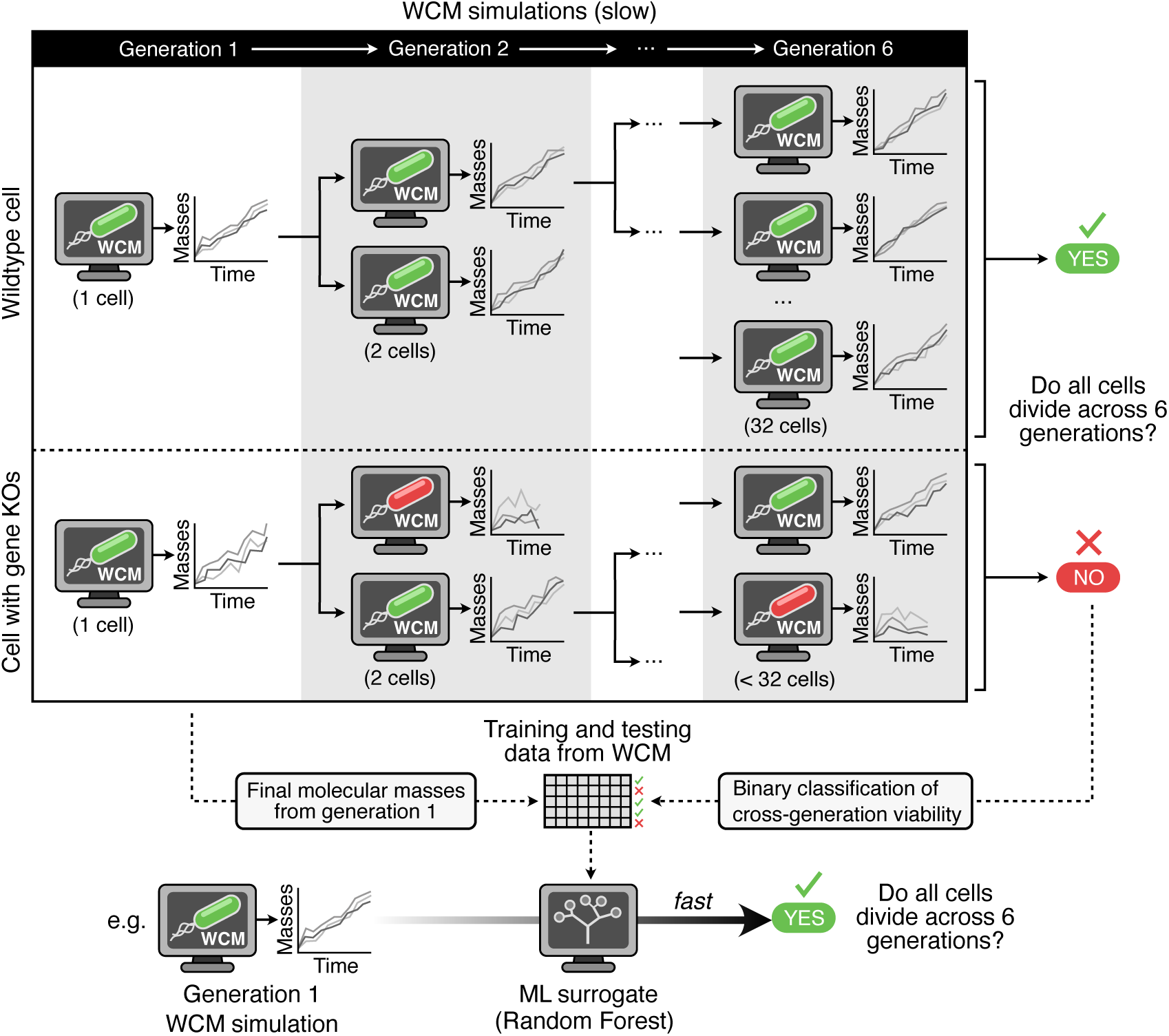
An example of a subset of the outputs of the *E. coli* WCM and the framework used to train a machine learning classifier that emulates the output of the WCM. Simulations start from two cells (a wildtype and a mutated cell) that are simulated for six generations. A random forest model is trained to emulate the behavior of the *E. coli* WCM (from generations 2-6). Data from the WCM output is used to train and test the machine learning surrogate. We use the value of the macromolecular masses at the end of generation one to train a random forest model that predicts whether all the cells in six generations will divide (e.g. the wildtype cell and all its replicates) or not (e.g. for the mutated cell with gene KO the red replicate in generation 2 did not divide). Once the model is trained, for future use, only one generation of the WCM will have to be run (instead of six), and the ML surrogate can be used to predict whether all the cells in six generations divide. This process is much faster.

Despite the richness of information enabled by the precise temporal tracking of cellular pro-cesses and components, this detail also imparts a substantial computational burden. As WCMs are developed for larger and more complex organisms, their computational complexity can increase substantially. To address this, when scaling up from *M. genitalium* to *E. coli*, the authors of the model implemented several computational improvements. These enabled the simulation run-time to decrease by two orders of magnitude, from approximately 10 hours for the *M. genitalium* WCM to 15 minutes for the *E. coli* WCM. This was achieved by writing inner loops in Cython and C, warm-starting the linear solver of the metabolic model and improving the file input and output (I/O).^3^ This boost in execution speed together with the substantial amount of *E. coli* data in the literature, enabled the modeling of cell division and growth such that several generations of cells could be simulated.^3^ This suggests that despite simulation of the *E. coli* WCM being much faster than for the *M. genitalium* model, the increase in the number of modeled genes and the ability of the model to simulate several cell divisions (i.e. generations) enlarges the possible combinations that have to be tested when using these models to predict phenotypes from genotypes.

The complexity of WCMs in general, stems from the massive number of time series they generate, tracking the tens of thousands of variables (i.e. molecule counts). However, in many applications, researchers are only interested in a specific subset of these outputs (e.g., the overall cellular mass). ML surrogates represent a solution to address this computational burden by exclusively making predictions for the outputs of interest after being trained on data generated by a WCM. ML surrogates, or emulators as they are sometimes known, are black-box models trained to approximate the behavior of complex and computationally expensive mathematical models.^15^ The major advantage of their use is the reduced computational resources required, allowing for the simulation time to fall from hours or days to seconds. ML surrogates are trained using data simulated by mechanistic models, taking as input their initial conditions and/or parameters, and they are designed to predict some or all of the original model’s outputs. The choice of ML algorithm depends on the nature of the problem, the amount and type of available data, and the desired accuracy and interpretability. ML surrogates have been used to emulate several biomedical mechanistic models^16–18^ and some molecular systems in biology,^19, 20^ allowing for efficient exploration of a mechanistic model’s parameter space,^21, 22^ as well as supporting model optimization^23^ and sensitivity analysis.^20^ ML surrogates have also been used to emulate mechanistic models based on Partial Differential Equations (PDEs).^16–18^ To date, models based on Ordinary Differential Equations (ODE) have only been used to show how ML surrogates can be trained, however no significant improvement in computational time was observed for systems analyzed to date.^19, 20^

In this study, we use a random forest-based ML surrogate trained on data from the single-cell *E. coli* WCM to estimate the viability of *E. coli* cells with reduced genomes (i.e. genomes containing fewer genes). Minimal genomes, containing only the essential genetic material necessary for cell reproduction, are generally used to help clarify the basic functions necessary for cell viability.^13^ Recently, there is also increasing interest in using them as simplified cellular chassis in synthetic biology to achieve more reliable and robust results.^13, 24–26^ Experimental studies have aimed to design reduced genomes for different organisms, removing as many genes as possible, while ensuring the cell remains viable (i.e. is able to divide and replicate).^27–29^ This process usually involves targeted design and gene selection based on experimental testing or biological knowledge.^11^ Since this is a time-consuming and laborious procedure, researchers often restrict their explorations to a few paths, building on existing literature. For instance, experimental attempts to engineer streamlined *E. coli* MG1655 cells able to grow in minimal media began in 2002 by removing 12 genome segments, with subsequent studies in 2006 and 2014 eliminating 31 and 13 additional segments, respectively.^30–32^ Further complexity in the design of reduced genomes comes from the interdependence between gene essentiality and the broader genomic context. A gene is considered essential if its removal from the genome affects the cell’s ability to divide and non-essential otherwise.^11, 33, 34^ However, some essential genes can become non-essential upon the removal of other genes. Similarly, some non-essential genes can become essential if another gene is removed or the growth media is changed (synthetic lethality).^11^ Predicting this type of interactions based on biological knowledge when hundreds of genes are deleted, is a significant challenge and highlights why experimentally focused projects that minimize genomes often leverage previous findings. Computational models offer a solution to these challenges by enabling the simulation of tens of thousands of gene deletions, uncovering new potential starting points for reduced genomes with a minimized risk of failure. For example, MinGenome represents an *in silico* approach based on a genome-scale metabolic model and mixed-integer linear programming to iteratively delete regulatory and metabolic genes in *E. coli*.^35^ Here, we propose an alternative approach that uses a WCM to design a reduced *E. coli* genome, allowing for the further deletion of non-metabolic genes and providing a more comprehensive representation of the genomic context and thus expanding the range of potential mutations compared with existing computational models.

WCMs have already been used to design minimal genomes *in silico* for *M. genitalium*.^11^ This was possible due to the small size of the genome (401 genes modeled) and the fact that the cells were simulated for only one generation.^11^ In contrast, the *E. coli* single-cell WCM used here captures the functionality of 1219 genes and can be run for as many generations as we choose.^3^ Given this additional complexity, we estimated that using a similar approach to the one used for *M. genitalium* would not be possible for *E. coli* without reducing the simulation time.^11^ To further illustrate this, consider that we want to simulate 10 genomes using the *M. genitalium* WCM. If there are 10 nodes available on a computing cluster, this will require approximately 10 hours because all 10 genomes can be run in parallel. If we want to simulate 10 genomes using the *E. coli* WCM for six generations (this study will later show why this is an optimal number of generations) and we assume that all cells in the six generations divide, we will need to simulate 630 cells in total (63 cells for each genome because at every division a cell produces two daughter cells). If we have 10 nodes available, simulating the 630 cells would take 15.75 hours (each cell simulation depends on the simulation of the mother cell, therefore, the generations need to be run sequentially and the number of cells grows exponentially with the generations). Given that the *E. coli* WCM contains three times more genes compared to the *M. genitalium* WCM, and the combinatorial aspect of gene deletions when searching for a reduced genome, simulating tens of thousands of genomes for six generations using the *E. coli* WCM is not feasible without using a surrogate model.

The aim of the current study is to show that the *E. coli* single-cell WCM can be used to design genomes with reduced size. From a methodological perspective, we show that this is possible with the help of ML surrogate models that can reduce the computational demand of a WCM and a genome design algorithm that aims to increase the number and size of segments deleted from the genome. From a biological perspective, we show that gene essentiality in the WCM matches *in vivo* gene essentiality from the literature, emphasizing that the search for a reduced *E. coli* genome is starting from data that has been empirically validated. We also perform a thorough comparison between our reduced genome and another reduced *E. coli* genome tested experimentally that grows in the same media and environment. Our analysis reveals some overlap between the two genomes, but it also emphasizes new deletions that are biologically sensible based on gene ontology (GO) analysis, highlighting the advantage of computational methods in designing reduced genomes. Furthermore, we show that the reduced genome obtained using the WCM is also able to delete non-metabolic genes, something that other *in silico* methods were not able to do. To the best of our knowledge, the present study is the first to show how an ML surrogate can emulate a large and complex biological model that integrates different mathematical modeling techniques and contains over 10,000 equations. Furthermore, this represents the first example of how the *E. coli* WCM can be used for synthetic biology applications.

## Results

### Data preparation for the ML surrogate

The ML surrogate presented in this study is trained on data produced by the *E. coli* single-cell WCM and aims to predict whether a reduced genome produces viable cells that successfully divide for several generations. Before we could train the ML surrogate, it was necessary to perform some data preprocessing on the WCM outputs. Firstly, we needed to decide how long the simulations should be run for (e.g., the number of generations) and what outputs/states from the WCM simulations would be used to train the ML surrogate. Preliminary *in silico* experiments representing single gene knock outs, where we simulated cell division for four generations showed that in some cases all the cells divided but also exhibited significant growth defects, especially in protein mass (Figures S1 and S2). This suggested that more than four generations should be simulated in order to accurately estimate the viability of a genome. We ran simulations for eight generations of those knock outs for which all cells divided in four generations but suffered growth defects and observed that 97% of the genomes that failed to divide in eight generations, did so in the first six generations(Supplementary Text, Figure S3). Therefore this number of cell divisions was used to generate the ML surrogate training data. Figure S3 shows a histogram of the generation when the first cell death occurs in simulations of eight cellular divisions representing single gene knock outs.

Additional decisions had to be made regarding the features to be used. Given the high dimensionality of the WCM, we focused our analysis on the 22 different macromolecular masses produced by the WCM. These masses, such as the total mass of all protein molecules, RNA molecules, DNA molecules, or membrane mass (see the description of Table S1 for the full list), offer a concise summary of the cell’s state.^36^ Collectively, they provide insights into growth and division, metabolic activity, genetic stability, and environmental interactions. We compared the accuracy of different ML surrogates using the values of these variables (i.e. features) either at the beginning of the simulations (*t* = 0, generation 1) or at the end of generation 1, just before the first cell divides. Results showed that the three algorithms tested (random forest, XGBoost and K-Nearest Neighbors) performed similarly, with random forest showing slightly higher accuracy irrespective of the features used (Table S1). We saw an improvement of approximately 17% in the F1-score of the training and test sets when using values at the last time step of generation 1 as features. Therefore, for the rest of this study, we chose to use as features the state of the cell at the last time step before the first division. A summary of the training procedure that will be used in this study can be seen in Figure 1 and the data preparation techniques are summarized in the Supplementary Text. An overview of the simulations run to train the ML surrogates is presented in Table S2.

Upon further investigation, we found that many of the macromolecular masses used as features to train the ML model were highly linearly correlated, suggesting that not all 22 features were necessary. Figure S4 shows the correlation matrix between the 22 features. The Pearson correlation coefficient values shown in the matrix were used to detect pairs of features with an absolute Pearson correlation > 0.95. We then kept only one of these, allowing us to reduce the number of features used to train the ML model to 12 (cell mass, growth, DNA mass, tRNA mass, extracellular mass, protein mass, projection mass, pilus mass, mRNA mass, small molecule mass, instantaneous growth rate, membrane mass). More details on how this reduction was performed can be found in the Supplementary Text. This reduction did not affect the accuracy of the ML surrogate.

### Training and testing the ML surrogate

The ML surrogate was trained and tested using simulated data produced by the WCM. We ran 924 single gene knock out simulations, corresponding to the knock out of 462 individual genes, each run twice. To make sure that the training and test data were kept separate, 738 of these knock outs (corresponding to 369 unique genes, where the removal of each gene was simulated twice) were used to train the ML surrogate and 186 (corresponding to other 93 unique genes, where the removal of each gene was simulated twice) were used to test it (Table SE1). The data was split such that no single gene knock out belonged to both the training and test set. Gene knock out simulations were chosen to train the ML surrogate because simulating different gene knock outs represents a source of variability, as the cell has to adapt to a different expression level of one gene. Furthermore, the fact that each simulation was repeated twice means that stochastic effects were considered, since stochasticity is embedded within the WCM.

The 12 key macromolecular masses (identified following the correlation analysis) at the last time point of the first generation were then used by a random forest classifier to predict whether all cells in six generations divide (Figure 1). A random forest classifier was chosen due to its superior performance (Table S1), simplicity, explainability and robustness.^37^ One of the primary strengths of random forest classifiers stems from their high accuracy, achieved by aggregating the predictions of multiple decision trees that reduces the risk of overfitting compared to considering only a single tree. This ensemble approach allows random forests to handle complex, non-linear relationships between features and the target variable effectively. Random forest is also highly robust and capable of managing outliers, missing values, and high-dimensional datasets without requiring extensive preprocessing like feature scaling. Additionally, random forest provides insights into feature importance, which can be valuable for understanding data and guiding feature selection.^38^ To assess the random forest model proposed here we used *k*-fold cross-validation on the training set to tune the model’s parameters and avoid overfitting. We also tested the model’s performance on a hold-out test set (containing data not used when training). We obtained an F1-score of 0.92 on the training set, 0.9 when averaging the cross-validation results, and 0.93 on the hold-out test set (Table S3). The corresponding confusion matrices show that when the ML model predicts that not all cells in 6 generations will divide (i.e. the gene knock out impacted viability), it is correct for 259 out of 261 cases (99%) in the training set, and 74 out of 76 (98%) in the test set (Figure 2A). In the context of minimal genome design, this suggests that by using the ML surrogate it should be possible to limit the removal of essential genes that upon deletion from the genome cause failure in the division of at least one cell in six generations.

**Figure 2:**
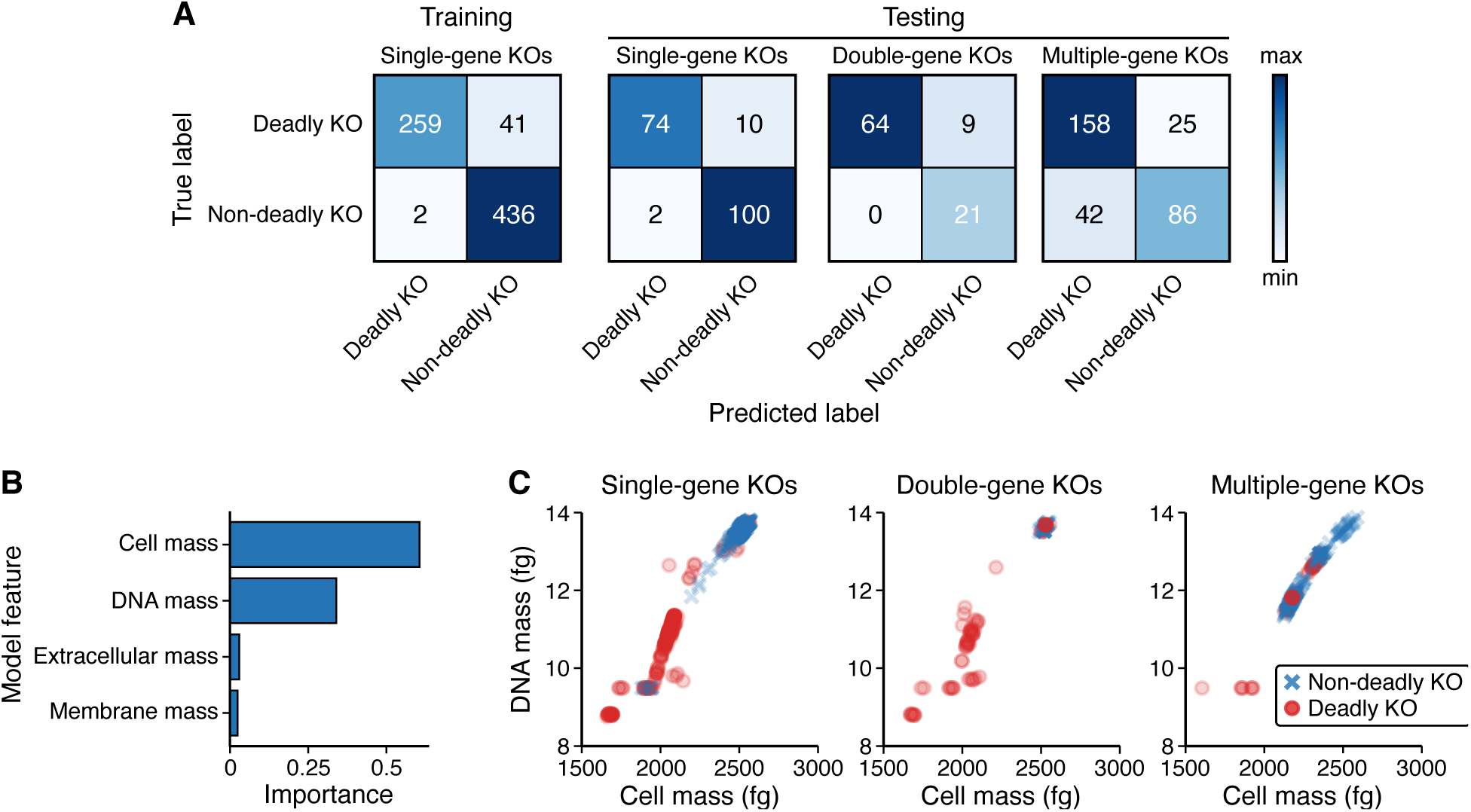
Performance of the initial ML surrogate trained on single gene knock out (KO) data. (**A**) The confusion matrices for the training of the ML surrogate on single gene knock outs, and testing on single, double and multiple gene knock outs. The true label is based on the output of the WCM and the predicted label is represented by the output of the ML surrogate. (**B**) The feature importance as outputted by the random forest algorithm trained on single gene knock outs. (**C**) Scatter plots of the two main features used for learning show different data sets containing single, double and multiple gene knock outs.

Our random forest classifier was trained, validated and tested repeatedly on these selected 12 features. We found that only four of the features (cell mass, DNA masss, extracellular mass and membrane mass) were necessary for making an accurate prediction. One of the advantages of using a random forest classifier is that it also ranks the importance of the features used for a prediction. Figure 2B shows the ranking of the features used by our classifier. More details about the impact of the features on the classifier and the misclassified data can be found in the Supplementary Text. This initial random forest surrogate was trained on single gene knock outs and tested on both single and double gene knock outs. The F1-score when testing the surrogate on double gene knock outs was 0.93 and there were no false positives (Figure 2A, Table S3). However, testing this initial ML surrogate on multiple gene knock outs (necessary for generating a reduced genome) resulted in an F1-score of 0.82 (Figure 2A, Table S3), a drop in performance when compared to the single and double gene knock out results. Figure 2C shows a scatter plot of the main two features used by the classifiers (at the end of generation one, just before cell division). We observed that when deleting more genes, the cell and DNA masses become similar among the deadly and non-deadly knock outs. Specifically, both the DNA and cell mass of the deadly knock outs are higher compared to when we knock out single genes. This suggests that in order to classify multiple gene knock outs, the surrogate has to be trained again using more features and including data representing multiple gene knock outs.

To address this, a new surrogate was trained by extending the initial training set of 738 single gene knock outs to additionally include 248 multiple gene knock outs, each run once for 6 generations (a list of the genes knocked out can be found in Table SE2). The F1-score of this model when tested on single and double gene knock outs remained similar, but when tested on the hold-out set of multiple gene knock outs (that were not used for training) it was 0.88, representing a 0.06 increase in F1-score (Table S4, Figure 3A) despite dealing with a much more challenging task compared to classifying single gene knock outs. When trained using the multiple gene knock outs, we used 11 of the uncorrelated macromolecular masses obtained following the correlation analysis described previously (the instantaneous growth rate was removed due to its similarity to the growth output perceived by the random forest). Figure 3B shows that the DNA mass, the membrane mass and the projection mass are essential for making accurate predictions. More information on the data used to train the new ML surrogate can be found in the Materials and Methods section.

**Figure 3:**
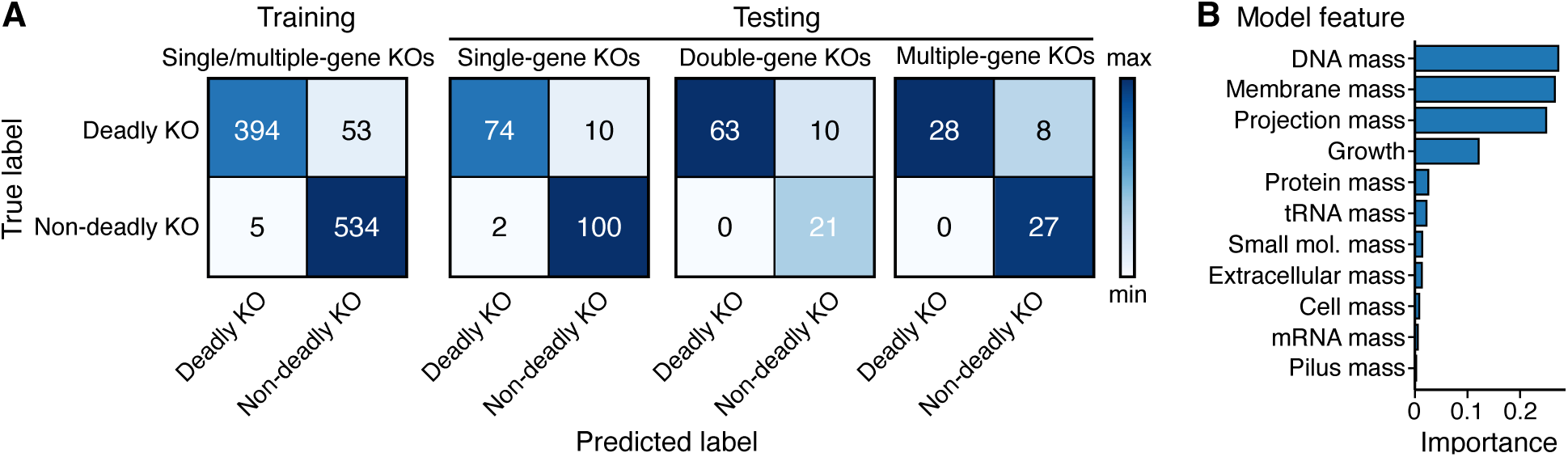
Results of the updated ML surrogate trained on single and multiple gene knock outs data. (**A**) The confusion matrices for the training and testing of the ML surrogate on single, double and multiple gene knock outs. The true label is based on the output of the WCM and the predicted label is represented by the output of the ML surrogate. (**B**) The feature importance as outputted by the random forest algorithm trained on single and multiple gene knock outs.

### Genome reduction

To demonstrate the value of the trained surrogate model, we showed that it can be used to design a reduced *E. coli* genome that still functions (i.e. divides) *in silico*. To do this, we used Minesweeper, an algorithm inspired by laboratory fragment-cassette-fraction genome engineering^29^ that employs a divide-and-conquer approach, which was used previously to find minimal genomes using the *M. genitalium* WCM.^11^ Minesweeper’s main aim is to generate smaller genomes quickly in a manner that is practical for *in vivo* testing. To achieve this, the algorithm splits the problem in smaller subproblems, solves each of them and then combines the solutions to address the bigger problem. The algorithm represents the design step in a design (identify possible gene deletions), simulate (the genome with those deletions) and test (analyze the simulated cell) cycle.^39^ Minesweeper requires simulating single gene knock outs and thousands of multiple gene knock outs. To be able to do this, we used our trained surrogate models.

Figure 4 shows the framework used to obtain the reduced genome using Minesweeper,^11^ which was employed to identify genes that can be removed from the genome, without losing the cell’s capability to divide successfully over six generations *in silico*. Minesweeper is given as input a list of genes that can be deleted (e.g. all genes in the WCM) and it outputs a run script to be used by a mathematical model that can estimate the essentiality of every gene in this list. Essentiality is estimated using the WCM and the initial ML surrogate. The values of the cell mass, DNA mass, extracellular mass and membrane mass at the end of the first generation of WCM single gene knock outs simulations are inputted into the random forest model that predicts whether for each genome, all cells in the first 6 generations would have divided. Minesweeper will only consider genes labeled as non-essential for future larger-scale deletions. This approach aims to maximize the design of viable reduced genomes early in the search process, while minimizing the potential negative effects of deleting essential genes.The initial random forest surrogate predicted that 1047 genes were non-essential and 172 genes were essential (Table SE3). Only the 1047 genes predicted as non-essential were considered for future deletions.

**Figure 4:**
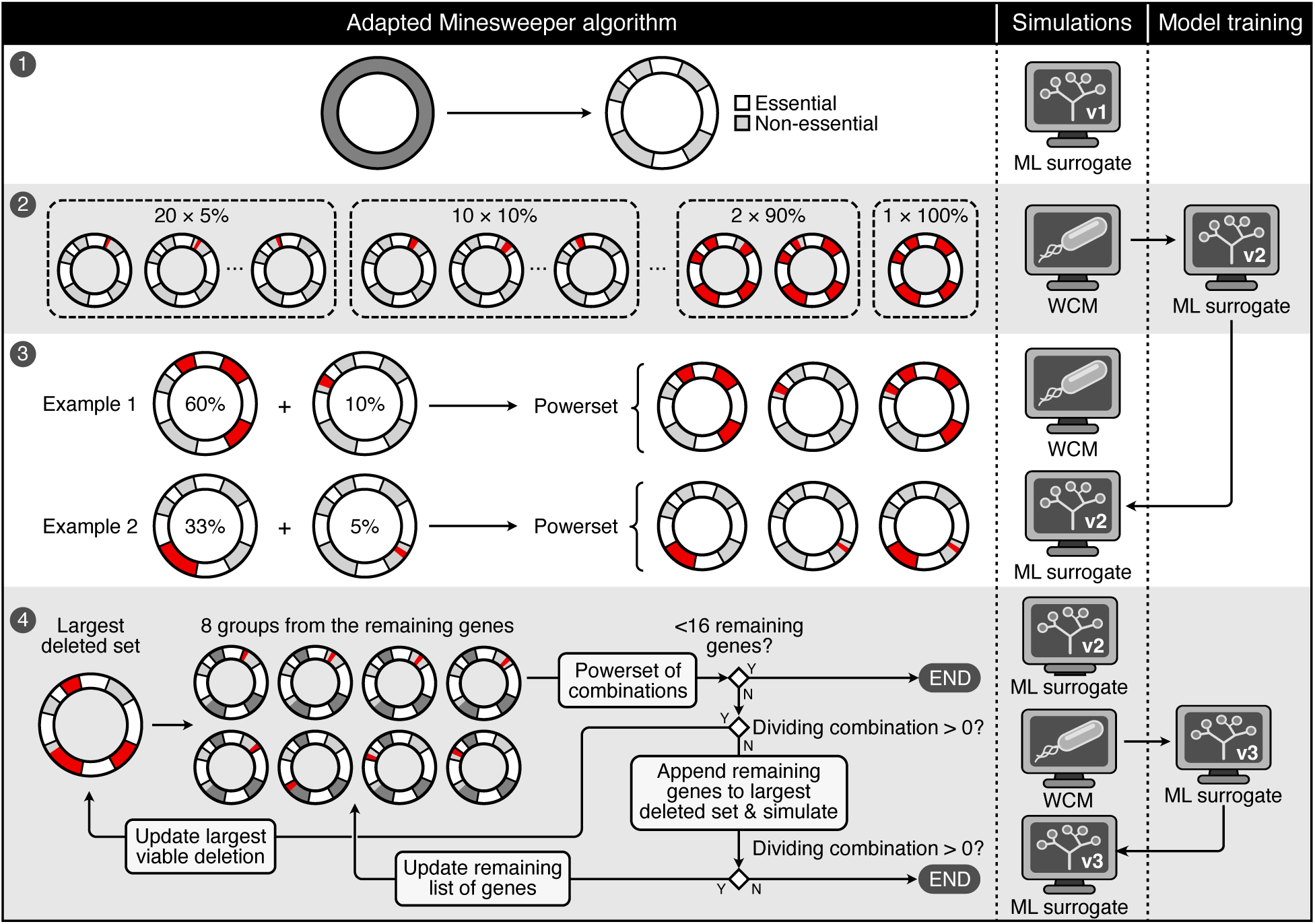
Summary of the framework used to generate the reduced genome. This is an adaptation of the Minesweeper framework,^11^ using the *E. coli* WCM^3^ and the ML surrogate. The process started by using the ML surrogate trained only on single gene knock out data (ML surrogate v1) to predict the essentiality of every gene. Only the genes characterized as non-essential go to stage two. In stage two, 56 groups representing different percentages of the non-essential genes were deleted individually from the genome (in red), and the corresponding simulations were run for 6 generations using the WCM. Permutations of these groups were used to train the new ML surrogate (ML surrogate v2). In stage three, we show examples where we took the two largest deletion segments that when removed produced cells that divided (example 1 and 2), appended a division-producing non-overlapping segments from stage two and created a powerset of combinations from these segments. . The fourth stage is cyclical and predictions were made using the newly trained ML surrogate (ML surrogate v2), randomly testing the dividing genomes using the six generations WCM simulations and training a new model with simulations from this stage (ML surrogate v3).

To provide initial experimental validation of the reduced genome we aim to obtain, we evaluated the predicted essentiality of genes using the initial ML surrogate (trained on data coming from the WCM) against data from a published genome-wide growth screen of *E. coli*.^40^ Our findings showed a strong agreement between the simulated essentiality and empirically defined essential genes. When viewed as a binary classification task, we designated non-essential genes as the positive (i.e., predictive target) class, given that Minesweeper only considers the removal of non-essential genes in its search for a reduced genome. Under these criteria, the *in silico* essentiality matches *in vivo* essentiality with an accuracy of 0.87 and an F1 score of 0.92 across all of the genes modeled by the WCM (Table S5). Furthermore, 94.3% of the genes labeled as non-essential *in vivo*, were also labeled non-essential by the ML surrogate (trained on data coming from the WCM). This single-gene knock out analysis demonstrates that the WCM and the ML surrogate adequately capture fundamental viability constraints, supporting the reliability of the non-essential gene identification during this initial step of our reduced genome design strategy.

Next, Minesweeper employs a top-down genome reduction strategy, where it aims to delete as many large segments of non-essential genes as possible.^11, 41^ The aim of the second stage is to identify gene segments of different sizes that could be removed without impacting division. This approach considers the proximity of genes, which is crucial for experimental deletions and aligns with the laboratory fragment-cassette-fraction genome engineering methodology.^29^ To begin with, the non-essential genes identified earlier are split into segments of different sizes, preserving their order. The second stage in Figure 4 outlines a subset of these segments. First, the non-essential genes are split into 20 segments, each containing 5% of consecutive non-essential genes. Similarly, the non-essential genes are also split into 10 segments of 10%, eight segments of 12.5%, four segments of 25%, three segments of 33% and two segments of 50%. For the larger segments (i.e. of 60%, 70%, 80% and 90%), Minesweeper considers the first x% and the last x% of non-essential genes (See the 90% example in Figure 4). In total, 56 segments of consecutive non-essential genes are formed. The number of deletion segments simulated represents the first modification made to the original version of Minesweeper.^11^ In the original version, the smallest simulated deletants included 12.5% of the non-essential genes. However, in our case these segments were the largest that when deleted, still allowed the cell to remain viable. This suggested that in order to minimize the cell further, we needed to simulate smaller deletion segments that can be combined in the next steps of Minesweeper. Therefore, we decided to adapt Minesweeper and also simulate the deletion of segments containing 10% and 5% of the non-essential genes, respectively.

In stage two, the full WCM was used to simulate the deletion of each of the 56 segments. From these, the removal of 20 of them produced dividing cells in six generations. These 20 segments correspond to two 12.5% groups, four 10% groups and fourteen 5% groups (Supplementary Table SE4). At this point a new ML surrogate was trained using combinations of these 20 segments in addition to the single gene knock outs used before. This new ML surrogate corresponds to the one described in the previous section (with performance presented in Figure 3) and to ML surrogate v2 in Figure 4. More details on the data used to train the surrogate can be found in the Materials and Methods section.

To minimize the genome even further without compromising division, Minesweeper takes the three largest segments that the WCM suggested can be deleted in stage two (two 12.5% groups and one 10% group) and for each of them it finds all non-overlaping segments of non-essential genes (from stage two) and creates powersets between these. The new ML surrogate trained on single and multiple gene knock outs was used at this stage to predict whether all cells in six generations divided when these combinations were deleted. To make sure that the genome representing the largest deletion that produced a dividing cell was carried to the next stage of Minesweeper, we tested the top 20 largest deletions using the full six generations simulations of the WCM. The main reason why this stage starts from the three largest deleted segments (instead of one segment) is because we wanted to have a wider range of starting deletions across the genome. The possibility of performing such an exhaustive search, combining the deletion of different non-essential segments, represents an advantage of *in silico* reduced genome design algorithms.

Up to this point, Minesweeper had identified the largest segments of non-essential genes that can be removed from the cell without preventing cell division and had combined these with other non-overlapping segments (of 12.5%, 10% or 5%, since these are the only ones that when deleted the cell remained viable) that can also be deleted according to the simulations run in stage two. To minimize the genome further, starting from the deletions already made, it would be necessary to append smaller segments than the 5% ones tried in stages two and three. Therefore, in stage four, Minesweeper started by taking the largest segment from stage three (for each of the three segments tested), which when removed produced a viable cell, and generated eight groups from the remaining genes, not included in the segment. The number of groups was determined by computational constraints when Minesweeper was first published.^11^ Following this, it created a powerset of combinations between these groups of genes and the current largest deleted segment. If some deletants formed by the removal of these combinations divided, it updated its record of the largest deletion segment and the remaining genes accordingly. If none of the combinations divided, it appended individual genes to the previous largest deletion segment (suggesting that the groups formed by the remaining genes are too large). If some of these divided, it updated the list of remaining genes. If none divided, the algorithm stopped. The algorithm also stopped if there were less than 16 genes after a powerset of group combinations is run. This means that if it were to continue, in the next iteration of the cycle, there would be 2 genes per group. In the original version of the Minesweeper algorithm, this limit was set to less than eight remaining genes (instead of 16). We modified the limit due to the higher number of modeled genes in *E. coli* compared to *M. genitalium*, which would lead to running a lot more iteration of this stage of Minesweeper. We ran this cycle three times starting from the largest deletions of each of the three groups from stage three. Minesweeper was stopped before converging due to the high number of simulations run (over 39,000) and the significance reduction in the genome’s size that had already been obtained. The newly trained ML surrogate was used to make predictions for these iterations, after which additional data was added to the training set and the surrogate was trained once again (see the Materials and Methods section). These three cycles produced three reduced cells (i.e. ignoring the single gene additions to the largest deleted set from stage four in Figure 4). The size of these reduced cells varies between 803 genes at the beginning of stage four simulations and 737 genes at the end.

The final and smallest reduced genome, EMine-737, contained 737 genes out of the 1219 modeled by the WCM (Table SE5). This represents a 40% reduction of the WCM’s genome in minimal M9 media supplemented with 0.4% glucose. To obtain this result, we simulated 1525 genomes running the WCM for six generations and 39,086 genomes running the WCM for one generation and predicting division using the ML surrogates. Assuming that we are running simulations using one CPU on a standard desktop computer and that all cells in six generations would have divided, running all simulations (1525 for six generations and 39,086 for one generation) would have taken approximately two and a half years. Without the ML surrogate (running all simulations for six generations) would have taken approximately 36 years, resulting in a reduction in computational time of 94% (14.4-fold reduction). The simulations used in this study were run in parallel, on a supercomputer, where possible.

### Reduced *in silico* genome (EMine-737) growth analysis

To further assess the performance of the ML surrogate and validate the viability of EMine-737, we simulated it 50 times for six generations using the original WCM. The results showed that for 96% of repetitions, all cells in six generations divide. We compared the growth and masses of one EMine-737 genome to those of one wildtype genome, where both were allowed to divide for six generations and produce two daughter cells at each division (Figure S8). We found that the median cell mass of the EMine-737 cells was 1309 fg compared to 1571 fg for wildtype cells.

The reduced genome cells also grow more slowly (0.14 fg/s median growth rate of EMine-737 compared to 0.22 fg/s for wildtype cells) and therefore take longer to divide compared to their wildtype counterpart. This may be due to the fact that the cells simulated by the WCM need to reach a certain mass before they divide. Figure S8 suggests that the RNA and protein mass of the reduced genome cells are often higher than those of wildtype cells (with a median protein mass of 232 fg for EMine-737 cells and 214 fg for the wildtype cells and a RNA mass of 104.6 fg for EMine-737 and 73.9 fg for wildtype cells).

To further assess the stability of the EMine-737 genome, we also ran simulations over 10 generation for 50 repeats (seeds) using the WCM, where each dividing cell produced only one daughter (to ensure reasonable computational demands). The results show that for 90% of these, all cells in 10 generations divided, compared to 98% for wildtype *E. coli* simulations.

### Reduced *in silico* genome (EMine-737) literature comparison

The construction of the *E. coli* WCM involved the integration of data from three distinct laboratory strains: K-12 MG1655 (used to obtain most high-throughput data) , B/r (used for composition data) and BW25113.^3^ This approach was necessary due to an insufficient amount of data available from a singular strain. This multi-strain data integration strategy might potentially constrain the capacity for direct comparison with other reduced *E. coli* genomes in the literature.

A number of reviews have compared existing laboratory reductions of the *E. coli* genome (primarily in K-12 MG1655 and K-12 W3110 strains),^26, 42, 43^ but only three studies tested the reduced MG1655 genomes in minimal media.^30–32^ First, the MDS12 strain published in 2002 deleted 12 genomic segments from MG1655 (8.1% deletion) without impacting growth rates in minimal media.^30^ A study published in 2006 removed additional regions from the genome to obtain the MDS43 strain (15.3% of the genome was removed).^31^ Later, another 13 genomic regions were removed to obtain the MS56 strain with 23% of the original genome deleted.^32^ Since MS56 contained the gene deletions from MDS12 and MDS43, we will compare EMine-737 to MS56.

Figure 5A shows a comparison between the genes of EMine-737, the genes included in the *E. coli* WCM and the genes included in MS56. This shows that the genomic regions deleted to obtain EMine-737 are different to the ones deleted in MS56. In fact, 79% of the genes deleted in EMine-737, have not been deleted in MS56. Figure 5B shows the proportion of genes included in each of the three genomes with respect to the total number of genes present in *E. coli* MG1655. Given that the version of the *E. coli* WCM used in this study includes the functionality of only a fraction of the total number of genes, it is unlikely that the EMine-737 will also be viable *in vivo*. However, a subset of the genomic regions removed to obtain EMine-737 could represent a new starting points for deriving minimal genomes *in vivo*; similar to the way MDS12 represented a starting point for deriving MDS43 and MS56.^30–32^ A list of the genes deleted in MS56, the WCM genes, and the genes deleted in EMine-737, are presented in Table SE6.

**Figure 5:**
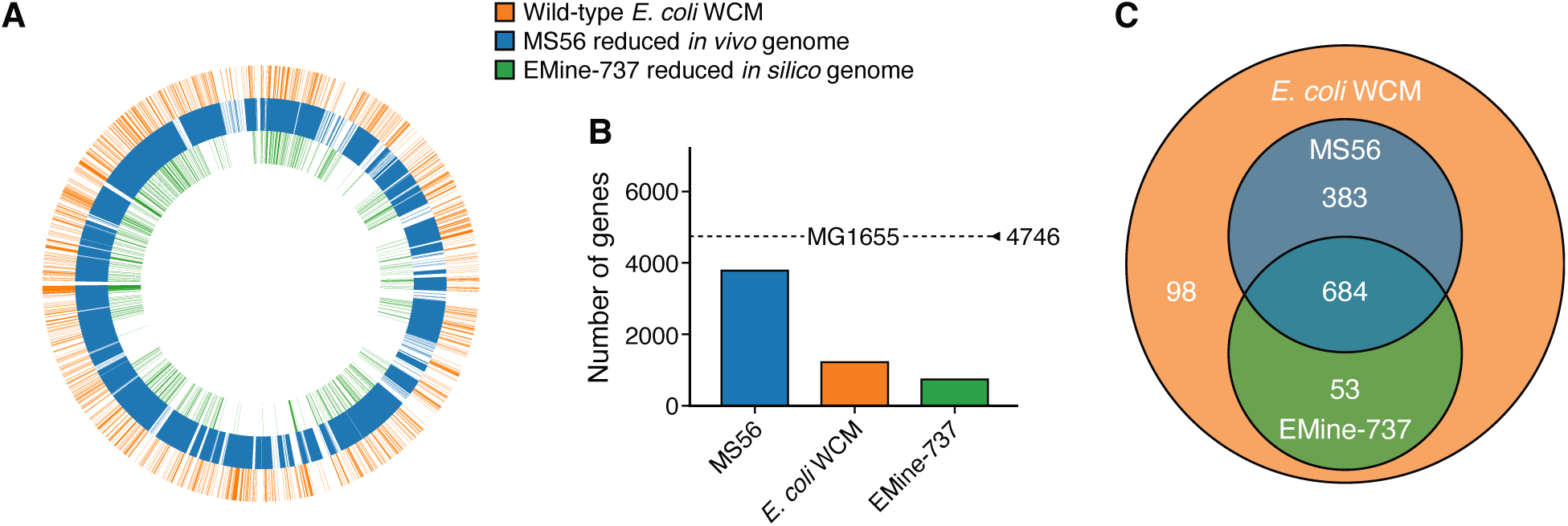
A comparison between the genome of EMine-737, the *E. coli* WCM wildtype genome and the MS56 genome. (**A**) Genes included in the *E. coli* WCM (the outer circle in orange), the genes included in MS56^32^ (the middle circle in blue) and the genes included in EMine-737 (the inner circle in green). (**B**) Proportion of genes included in each of the genomes relative to a wildtype MG1655 genome. (**C**) Number of genes included in each genome from the total number of genes included in the *E. coli* WCM. The numbers sum up to 1218 because gene *ilvG* is split in two genes in the WCM.

To assess the advantage of using a WCM for designing a minimal cell, compared to genome scale metabolic models, we calculated the number of non-metabolic genes that were removed from EMine-737. In total there are 89 genes that were removed from EMine-737 that are not modeled by the genome scale metabolic model iJO1366 used in MinGenome.^35, 44^ These genes are also listed in Table SE6.

### Gene ontology analysis comparing EMine-737 to MS56

To better understand the relationship between EMine-737 and MS56, we performed gene ontology (GO) analysis for the 53 genes included in EMine-737 but not in MS56, the 684 genes included in both, the 383 genes included only in MS56, and the 98 genes not included in either (Figure 5C). In all cases we analyzed the GO terms for which all genes were removed, unless otherwise stated.

### Genes present in both EMine-737 and MS56

Table SE7 contains the genes and GO terms for the 684 genes (92.8% of the genes in EMine-737) included in both EMine-737 and MS56. The GO terms for which all genes are present in both genomes are also represented visually in Figure 6A. Further analysis of these genes showed that essential processes involved in ribosomal function (e.g. small ribosomal subunit, large ribosomal subunit, large ribosomal subunit rRNA binding) were present in both reduced genomes. In addition, we found that both reduced cells are able to breakdown and synthesize amino acids and intermediates, since GO terms involving the catabolism of glycine, D-arabinose, L-arabinose, and arginine as well as the biosynthesis of tyrosine, diaminopimelate, homoserine, lysine and glutamate were present in the two genomes. This suggests that the cells maintain the molecular machinery responsible for protein synthesis. Important energy production mechanisms are also present in both genomes, which is supported by the retention of genes involved in oxidative phosphorylation, proton-transporting ATP synthase complex, succinate dehydrogenase complex and glycerol-3-phosphate-transporting ATPase complex. Genes involved in the bacteria’s ability to import nutrients from the environment also remain (e.g glucose import across plasma membrane, ATP-binding cassette (ABC) and methionine-importing ABC transporter complex), as well as genes responsible for the transport of ions and small molecules (e.g. spermidine, D-methionine, Vitamin B6 Transport, Methionine and Glucose Import Across Plasma Membrane, Mannose Transmembrane Transport). Other components that are important for the function and structure of cellular machinery, such as nucleotide synthesis and metabolism, are also maintained in both reduced genomes.

**Figure 6:**
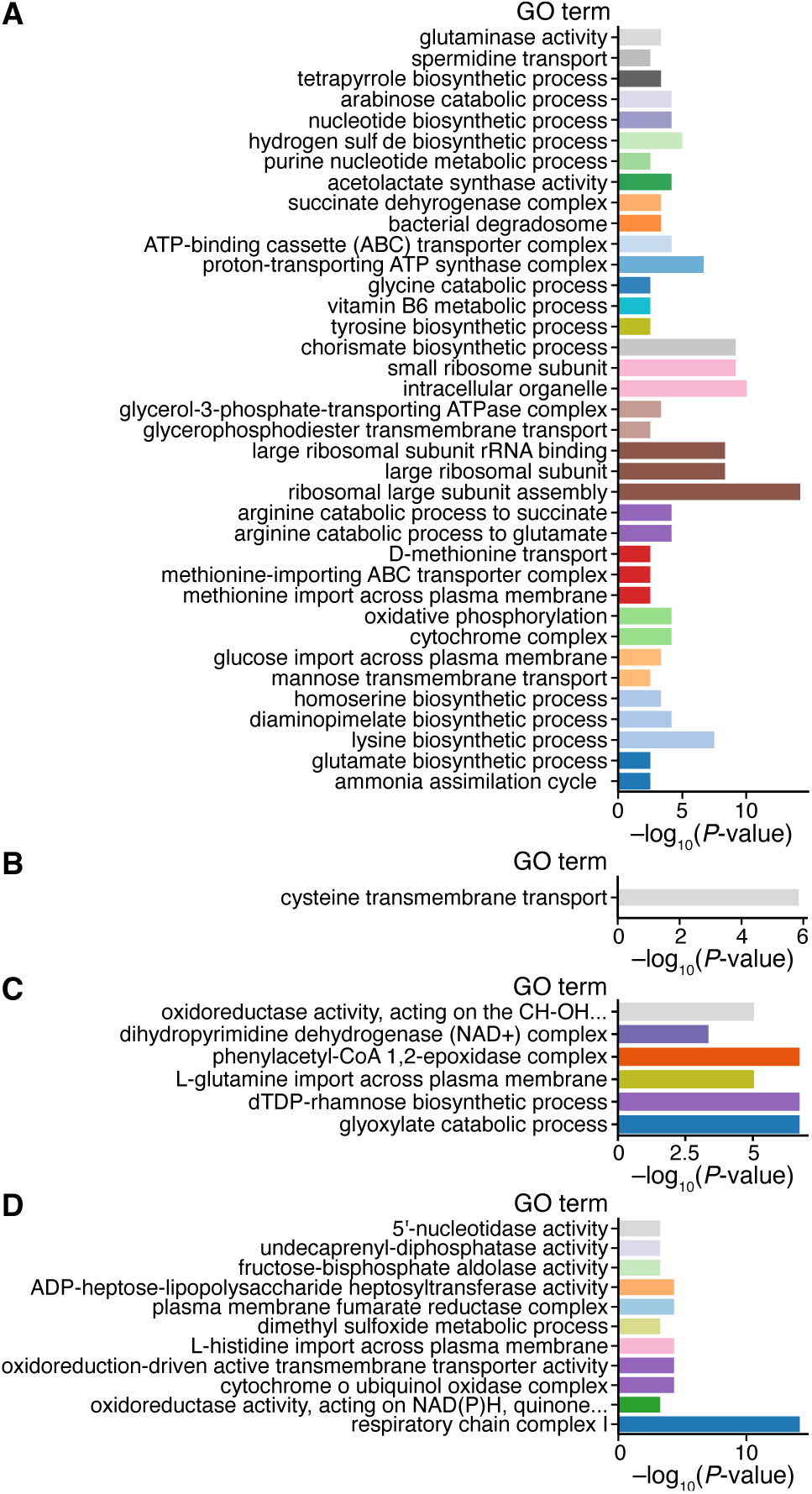
GO term analysis of reduced genomes. **(A)** The GO terms for which all genes are present in both EMine-737 and MS56. Terms that are next to each other and are represented using the same color have genes in common. **(B)** The GO terms for which all genes were deleted in MS56, but are present in EMine-737. **(C)** The GO terms for which all genes were deleted from both MS56 and EMine-737. **(D)** The GO terms for which all genes were deleted from EMine-737, but are present in MS56. Terms that are next to each other and are represented using the same color have genes in common. The p-value in all panels represents the probability that the observed enrichment of a GO term occurred by random chance.

### Genes present in EMine-737, but not in MS56

Further gene ontology analysis was performed for the genes that are kept in EMine-737, but were deleted in MS56. There are 53 such genes that correspond to 5 gene ontology terms (Table SE8) representing different categories of cysteine transport (with 100% of the genes removed, Figure 6B), L-cystine transport (with more than 75% of the genes in that ontology term removed), sulfur utilization (with 60% of the genes included in that ontology term removed) and lyase activity (with 7% of the genes included in that ontology term removed). In M9 media, the primary sulfur source is typically sulfate, not cystine.^45^ Therefore, the removal of *tcyJLN* (cystine ABC transporter) genes would have little direct impact on sulfur acquisition. The *dcyD* gene (encoding D-cysteine desulfhydrase) is mainly relevant when D-cysteine is used as a source of sulfur. Given that this is not the case in M9 media, its deletion shouldn’t impact the cell a lot.^46^ This analysis suggests that indeed these genes could be deleted, like they were in MS56. In fact, these four genes were among the remaining genes that would have been considered for deletion had we allowed Minesweeper to converge.

We simulated the EMine-737 reduced genome with the additional deletions of genes *tcyJLN* and *dcyD* for six generations (with two daughter cells for each division) and for 10 generations (with one daughter cell for each division), with each simulation repeated 20 times. For both sets of simulations, 18 out of the 20 repetitions produced viable cells in six and 10 generations. This suggests that indeed these genes can be deleted *in silico* as well.

Analyzing further the 53 genes that were removed in MS56, but not in EMine-737, we found that while MS56 removed all genes from the *als* operon, EMine-737 still contains three of them (*alsACK* encoding D-allose ABC transporter units and D-allose kinase). These three genes were kept in EMine-737 because in the fourth stage of Minesweeper, when they were individually appended to the largest deleted segment, the cell did not divide when these segments were deleted. This represents an example where the deletions from an operon were split because deleting the corresponding genes individually (in addition to the largest already deleted segment) did not produce a dividing a cell. The *lac* operon is another example of operon deleted in MS56, but not in EMine-737. The *lacA* (encoding galactoside O-acetyltransferase) and *lacZ* (encoding -galactosidase) genes are among the remaining genes that would have been considered for deletion if we did not stop Minesweeper before converging, while the deletion segment containing the *lacY* gene (encoding lactose permease) did not produce dividing cells in the fourth stage of Minesweeper.^46^ A similar behavior is observed for the *mhpDEF*, *phnCDEN* and *prpBCDE* genes. In the case of the *put* operon, when adding the *putA* (encoding a flavoprotein with functions such as transcriptional repressor and membrane-associated enzyme) and *putP* (encoding a sodium/proline symporter) genes to the largest deletion during the fourth stage of Minesweeper, we see that both deletions cause a failure in division for the respective genomes.^46^ The same pattern is observed for the *yagE* and *yagF* genes (encoding for different CP4-6 prophage enzymes). We tested the viability of EMine-737 with additional deletions of *tcyJLN, dcyD, alsACK, lacAYZ, mhpDEF, phnCDEN, prpBCDE, putAP* and *yagEF*, running 20 repetitions of six generations simulations (with each viable cell producing two daughter cells) and 10 generations simulations (with each viable cell producing one daughter cell). The results showed that out of the 20 repetitions, for the six generations simulations only 12 saw all the cells divide, while for the 10 generations simulations, 15 saw all the cells dividing. This represents an example of how data analysis from *in silico* and *in vivo* reduced genomes can be used to derive even smaller genomes.

### Genes removed from both EMine-737 and MS56

Table SE9 contains 98 genes (20% of the genes removed in EMine-737) that have been removed from both EMine-737 and MS56 and the corresponding gene ontology terms. There are six gene ontology terms that contain genes that are completely removed from both EMine-737 and MS56 (Figure 6C). Phenylacetyl-CoA 1,2-epoxidase complex is one of these. This complex is encoded by the *paaABCDE* genes and is involved in the phenylacetate degradation pathway under aerobic conditions.^47^ This process is important for *E. coli* when phenylacetate is used as a carbon and energy source. However, both in EMine-737 and MS56, the main carbon source is glucose, therefore the removal of this complex should not cause significant problems to the cell’s growth. The dTDP-rhamnose biosynthetic process is also impaired in both MS56 and EMine-737. The dTDP-rhamnose is a sugar nucleotide that is not directly involved in the metabolism of glucose, so its absence is unlikely to have an immediate impact on *E. coli*’s ability to utilize glucose as a carbon source for energy production.^46, 48^ Another term removed is glyoxylate catabolic process. Glyoxylate is an alternative to the tricarboxylic acid (TCA) cycle in *E. coli*. It allows the bacterium to grow on acetate or fatty acids as sole carbon sources, but with glucose available in the media it becomes less important.^49^ The process involving the import of L-glutamine across plasma membrane is also impaired in both reduced genomes. Although this might affect glutamine synthesis, further investigation showed that genes involved in the internal synthetization of L-glutamine (*glnA*) are still present in the EMine-737 genome and the reduced cell is producing glutamine (Figure S9). Other gene ontology terms removed from both EMine-737 and MS56 were the oxidoreductase activity, acting on the CH-OH group of donors, quinone or similar compound as acceptor and dihydropyrimidine dehydrogenase (NAD+) complex.

### Genes removed in EMine-737, but present in MS56

The GO terms for the 383 genes that were deleted in EMine-737, but kept in MS56 together with the genes themselves can be found in Table SE10. Figure 6D also shows the GO terms that for which all corresponding genes were removed in EMine-737, but present in MS56. Several ontology categories related to respiration and oxidase complexes (e.g., the cytochrome o ubiquinol oxidase complex and respiratory chain complex I) as well as oxidoreduction and oxidoreductase activities, were removed in EMine-737. Under aerobic conditions, *E. coli* can utilize alternative terminal oxidase pathways for energy production, such as the cytochrome *bd*-I and cytochrome *bd*-II complexes, instead of the cytochrome o ubiquinol oxidase complex and respiratory chain complex I.^46^ The genes involved in oxidoreduction transmembrane activity and cytochrome-c oxidase activity largely overlap with or are subsets of those involved in the cytochrome o ubiquinol oxidase complex. These deletions reflect a shift toward using alternative oxidases, simplifying the electron transport chain while maintaining aerobic respiration in a minimal media environment. Additionally, the *cyoABCD* genes, which encode components of the cytochrome o ubiquinol oxidase complex, are classified as non-essential in the literature.^46^ The dimethyl sulfoxide (DMSO) metabolic process is also removed from EMine-737, which represents a terminal electron acceptor in anaerobic conditions,^50^ but without significant activity in aerobic conditions. Similarly, the plasma membrane fumarate reductase enzyme complex removed in EMine-737 is catalyzing the reduction of fumarate to succinate in anaerobic conditions. However, its absence in aerobic conditions is unlikely to affect growth or function.^51^

Another set of deletions involves genes associated with the cell wall and outer membrane biosynthesis. The genes representing the activity of ADP-heptose-lipopolysaccharide heptosyltransferase ( *waaC, waaU, waaQ* and *waaF*) are involved in the biosynthesis of lipopolysaccharides (LPS), a region in the outer membrane. Mutants lacking the *waaC* and *waaF* genes have shown increased sensitivity to a variety of antibiotics, including β-lactams and aminoglycosides.^52^ This hypersensitivity is likely due to the compromised integrity of the bacterial cell envelope, which is essential for resisting external threats, but would probably be less problematic in a well-defined environment. The genes *ybjG, lpxT* and *pgpB* are involved in the metabolism of undecaprenyl pyrophosphate (Und-P), a lipid carrier for glycan biosynthetic intermediates.^53^ Deletions involving some of these genes have been shown to increase sensitivity to fosmidomycin, an antibiotic that inhibits Und-P biosynthesis.^53^

Three GO terms involved in nucleotide and carbohydrate metabolism are also fully removed in EMine-737 genome. These include the overlapping terms 5’-nucleotidase activity and XMP 5’-nucleotidase activity (not shown in Figure 6 due to the overlap in genes) and the corresponding genes *yrfG, yjjG* and *ushA* (encoding for different nucleotidase), which are involved in the metabolism of purine, in maintaining nucleotide homeostasis and ensuring balanced pools of nucleotides and sugars for cellular functions, particularly under stress or in nutrient-limited environments.^46^ Genes *fbaB, lsrF* and *fbaA* are also deleted in EMine-737. These are involved in glycolysis and gluconeogenesis, as well as in the fructose-biphosphate aldolase activity term. While mutants of *fbaB* and *lsrF* exhibit growth in several media, mutants of *fbaA* exhibit no growth in LB enriched, LB Lennox and Luria-Bertani broth and M9 minimal media.^46, 54^ Therefore, the deletion of this gene has to be investigated further before including it in any experimental deletions.^46^ The other gene ontology term removed from EMine-737, but not MS56 represents the import of L-histidine across plasma membrane. This process allows the uptake of L-histidine from the environment. Despite this process being impaired in EMine-737, *E. coli* is likely able to synthesize this amino acid internally. In fact, all the genes involved in the biosynthesis pathway of L-histidine (*hisABCDFGHI*) are still present in EMine-737 and the cell is producing the amino acid (Figure S9), although in smaller amounts compared to the wildtype cells (with a median L-histidine molecule counts of 52,483 per cell for EMine-737 and 75,704 per cell for wildtype).

The analysis above emphasizes the fact that the deletions observed in EMine-737 represent a distinct minimal genome design compared to previous literature-reported minimal *E. coli* genomes. The investigation on entirely removed GO terms from EMine-737 demonstrates that the deletions generated entirely by the WCM can be interpreted biologically and could guide future experimental design when testing the deletions proposed by EMine-737. The advantage of using the WCM stems from its ability to test the impact of novel combination of gene deletions on cell growth, energy balance and metabolic dynamics, giving a system-level insight. While some deletions can be identified and tested based on gene ontology analysis alone, assessing the impact of multiple deletions in combination requires either a mathematical model that represents the cell at a system-wide level or extensive experimental validation, which can be prohibitively expensive.

## Discussion

In this study, we demonstrated that employing an ML-based surrogate to replicate the viability predictions of an *E. coli* WCM enables the use of such models for synthetic biology applications, including the *in silico* search for reduced genomes. Specifically, we showed that a search on a typical desktop PC that would take over 36 years, can be reduced to approximately 2.5 years, representing a speed-up of around 14-fold. With access to a supercomputer that allows running up to 40 simulations in parallel, the equivalent simulation time using only the WCM is 666.2 days, and using the WCM together with the surrogate is 35 days, showing that the surrogate enables a 95% decrease in computational time (see the Materials and Methods section for more details). These results emphasize that even with the help of supercomputers, WCMs are prohibitively expensive and difficult to use in scenarios that require simulations of many genomes without the use of surrogate models. It is important to highlight that while simulation time is notably reduced via the use of a surrogate, we lose some of the detail and granularity of the WCM simulations. Specifically, by only using the WCM to simulate the dynamics of one cell, for one generation, and then using the ML surrogate to predict whether all cells in the future five generations divide, we lose any information about the dynamics of the cells from future generations. This could potentially overlook some delayed effects of gene deletions on cell viability and affect our understanding of underlying biological mechanisms such as the cell’s response to stress during those generations. Furthermore, using only a subset of the macromolecular masses from the end of the first generation as features to predict future divisions, the ML model is not able to capture the full complexity and interactions among all variables of the WCM. These features were enough to build a classifier that predicts whether all cells in six generations divide. However, for more complex tasks, such as the prediction of yield of a specific amino acid or metabolite, it would probably be necessary to use data regarding the dynamics and concentration of specific molecules in order to build an accurate ML surrogate. Other insights that might be lost by using ML surrogates are the subtle interactions between different molecules that are captured by simulating a dynamical model. To address this, we used the entire WCM simulations in instances where we were interested in studying the interactions and the dynamics of the cell at the molecular level. For example, we used the full WCM to investigate the dynamics of amino acids such as L-histidine and L-glutamine in our GO analysis section (Figure S9) and to confirm that the reduced genome we derived using the ML surrogate is indeed viable when simulated using the original WCM.

In this study, for designing reduced genomes, we used our ML surrogate model to predict a binary variable (cell division), but we are confident that this approach can be expanded to predict continuous variables such as metabolic yield, or to forecast time series corresponding to some variables of interest. To predict continuous variables, it would be necessary to train regression-based surrogate models, while for time series forecasting, deep learning approaches such as Long Short Term Memory (LSTM) networks could be used. These methods could be useful when using the *E. coli* WCM for metabolic engineering, particularly when many simulations need to be run.

Typically, ML surrogates are trained on initial conditions and/or the parameters of mechanistic models. In this study, this was not possible due to the high dimensionality and complexity of the WCM. We would have needed more than 10,000 training examples to avoid having more features than training data. Instead, the surrogate presented here was trained using feature variables that summarize the state of a cell (i.e. macromolecular masses). Moreover, rather than using the initial state of these variables, we chose to use the state at the end of the first generation of each genome simulation, which increased the F1-score of the classifier by 16-18%. This suggests that there is not enough information regarding the cellular dynamics at the beginning of each simulation to make an accurate prediction. Even with this approach, the number of cells that had to be simulated for each individual genome was reduced from 63 when running only the *E. coli* WCM for six generation to one using the ML surrogate (assuming that all cells in six generations divide).

The excellent performance of our surrogate, with a F1-score of 0.93 on single gene knock outs, suggests that the cell mass, DNA mass, extracellular mass and membrane mass at the end of generation one hold enough information to predict the future viability of daughter cells for another 5 generations. Regarding the false negative predictions made by the ML surrogate, they are likely caused either by the fact that the decrease in cell and DNA mass is not clear at the end of generation one, or because there is some other reason behind the failure in division that is not captured by the macromolecular masses. GO analysis performed on the misclassified genes (Table SE11), did not reveal any obvious pattern regarding a particular type of genes that are more likely to be misclassified. In addition, we analyzed the generations when the first cell fails to divide for all genomes in the training set of the initial ML surrogate (Figure S10A), the 41 genomes in the training set of the initial ML surrogate that were misclassified as deadly (Figure S10B), all genomes in the test set of the initial ML surrogate (Figure S10C), and the 10 genomes from the test set of the initial ML surrogate that were misclassified as deadly (Figure S10D). The results suggest that the ability of the ML surrogate to predict cell division is not influenced by the generation when the first cell fails to divide in a genome, since we can see in Figure S10B and D that the surrogate misclassifies more often when the failure in division occurs in generations one or two. In spite of the number of false negatives (in Figures 2 and 3), they do not represent a problem in the context of genome reductions, as misclassified knock outs will still be identified in later stages of reduction. In contrast, false positives (in Figures 2 and 3) are more problematic, as they decrease the potential pool of genes to be deleted when searching for a minimal genome. The ML surrogates presented in this study have a small false positive rate of 1-2%.

Initially the ML surrogate was trained on a set of individual single gene knock out simulations and tested on a separate set of individual gene knock out simulations, double gene knock out simulations and multiple gene knock out simulations. Figure 2 and Table S3 show that while this initial ML model was able to generalize well on novel single gene knock outs (not part of the training set) and double gene knock outs, it struggles to perform as well when tested on multiple gene knock outs. To address this limitation, a new ML surrogate was trained by adding to the original training set (that only contained single gene knock outs) data coming from multiple gene knock outs. Figure 3 and Table S4 show that the new model was able to generalize better to novel multiple gene knock outs, without losing its prediction capabilities for single and double gene knock outs. This suggests that the multiple gene knock outs data set allows the ML models to learn new dynamics. The lower performance of the ML models when tested on multiple gene knock outs can be caused by two factors. Firstly, it is possible that the macromolecular masses dynamics are different when multiple gene knock outs are performed and therefore adding more such data to the training set would be beneficial to learn the new dynamics (as shown in this study). Secondly, it is possible that some of the dynamics causing failures in division when multiple genes are knocked out, are not captured by the macromolecular masses. To address this, more features would need to be included when training the ML models such as, the distribution of fluxes in the metabolic network.

The *E. coli* genome obtained by using a combination of six generations of WCM simulations and the ML surrogates was reduced by 40%, from 1219 to 737 genes. A direct comparison to *in vivo* results is not possible because the WCM only includes the functionality of a fraction of *E. coli*’s MG1655 genome. However, the analysis performed in this work shows that 79% of the genes removed from the WCM to obtain EMine-737 represent novel deletions, that are not present in the experimentally verified MS56 reduced genome. GO terms analysis showed that the removal of some of these genes can be interpreted biologically given the specific growth conditions of the cells, and could guide future experimental design when testing the deletions proposed by EMine-737. The advantage of using the WCM stems from its ability to test the impact of novel combinations of gene deletions on cell growth, energy balance and metabolic dynamics, giving system-level insights. This becomes particularly important in the context of synthetic lethality (i.e. the unviability of multiple gene deletions, where individual deletions are viable). This often arises from non-obvious functional redundancies and distributed network buffering.^55^ These cases cannot be identified by discrete individual approaches such as GO analysis alone, and require a more holistic understanding and testing that can be guided by WCMs. Furthermore, unlike other mathematical models, WCMs are also able to simulate deletions of non-metabolic genes. This study has shown that 18.46% of the genes deleted in EMine-737 are non-metabolic and therefore have not been considered in previous *in silico* studies such as MinGenome.^35^

Mathematical models are not perfect and we are aware of the current limitations of the *E. coli* WCM. For example, not all functional genes are modeled yet, the WCM was built using data from several strains and it was not tuned to perform genome design tasks. These limitations can affect the reliability of the reduced genome we discovered. For example, we do not know the effect of unmodeled genes on growth and other phenotypes, especially when several genes are knocked out. Furthermore, while this study shows that the WCM is able to accurately predict the viability of single gene knock outs, no such test has been performed for multiple gene knock outs. For this reason, we propose that when the reduced genome presented in this study is tested *in vivo*, experiments should start by deleting the various 5% non-essential gene segments generated in the second stage of Minesweeper (found in Supplementary Table SE4). If needed, these segments could be divided even further to test the viability of even smaller ones. This would represent an initial step for testing the reduced genomes presented in this manuscript that would allow researchers to iteratively validate the performance of the WCM starting from fewer knock outs before building upwards. The genome analysis we performed also represents some proof that the deletions proposed by the WCM make sense biologically. Approximately 56% of the genes included in the WCM were kept both in EMine-737 and the reduced genome from the literature that we compared our predictions to (MS56), and 8% are removed from both. The 31.4% of genes that are removed from EMine-737, but kept in MS56 represent novel gene deletions arising from the study. The GO analysis performed suggests that indeed some of these genes could be removed under the environmental conditions tested in this work.

In this study, we used the Minesweeper algorithm to find reduced *E. coli* genomes. Due to the extensive size of the *E. coli* genome, we implemented two modifications to the algorithm. Firstly, in the second stage, which involves simulating the deletion of different segments of non-essential genes, we introduced smaller segment simulations, including those of 10% and 5% of the non-essential genes. This change was essential because initial results from the WCM indicated that the only viable deletions that did not inhibit cell division were two segments each of 12.5%. These findings suggested that without simulating smaller segments, Minesweeper would exhaust possible deletions too early, preventing further progress in subsequent stages. Secondly, in the fourth stage, we modified the stopping criterion of the algorithm to end when fewer than 16 genes remained, rather than the usual eight. This change was implemented to manage the computational demands, stopping the process early due to the high number of required simulations. Despite this modification, Minesweeper was stopped before converging due to the high number of simulations already run (over 39,000) and time constraints, emphasizing the importance of using a surrogate. As shown in the GO analysis, it is possible that if continued, Minesweeper would have led to a further reduced genome. As more genes are added to the *E. coli* WCM, the number of algorithmic iterations needed to search for reduced genomes will increase. As WCMs become more powerful and better at matching cellular phenotypes, their computational complexity needs to be addressed for genome design tasks, making the use of ML surrogates timely and advantageous.

Minesweeper is designed to function independently of specific computational models like the WCM or ML surrogates. It operates by processing inputs which include lists of genes for the initial stage and viability labels of previously simulated genomes for subsequent stages. Its primary outputs are configurations for new combinations of gene knock outs, along with runscripts tailored for the simulation model in use (in this case the *E. coli* WCM). This design underscores Minesweeper’s utility as a flexible, ’plug-and-play’ tool, adaptable to various genomic models beyond just WCMs or ML surrogates. For the specific objectives of this research, as mentioned above, we modified certain parameters of Minesweeper to better align with the size and increased complexity of the *E. coli* genome and WCM.

The inputs, outputs and parameters of Minesweeper do not depend directly on the viability measure used. In this work, we used the division of cells in six generation to define gene essentiality and genome viability. An alternative approach would be to use the growth rate of the cell. To illustrate how this work could be adapted for this, we trained an alternative ML surrogate, that uses the 12 uncorrelated macromolecular masses at the end of generation one to predict the average growth rate of all cells in generation four. The performance of this ML model on the same single gene knock outs used previously in the training and testing set (for the division classifier) is presented in the Supplementary Text and in Table S6. We observe that random forest again performs best, with an R-squared on the training set of 0.93 and on the test set of 0.89. This model can be extended to predict growth rate of multiple gene knock outs as well. To employ such model for designing reduced genomes using Minesweeper, this growth rate estimate will have to be transformed into a discrete variable. This could be done for example, by assuming that if the growth rate of a mutated cell is not significantly different to the growth rate of wildtype cells, then the genome being tested is viable. If the growth rate is significantly different to that of wildtype cells, the genome is not viable. This definition of viability can then be used directly within Minesweeper, by simply replacing the cell division categorization with this growth rate similarity categorization.

## Acknowledgments

We would like to thank the Advanced Computing Research Centre (ACRC) at the University of Bristol for access to the BlueCrystal and BluePeble supercomputers. Special thanks to the HPC and RDSF teams of the ACRC, particularly Dianaimh Greene and Callum Wright for their help with BlueCrystal and BluePebble. Furthermore, we would like to thank Oracle Cloud for allowing us to run some of the simulations used in this study on their clusters. Special thanks to Dr. Christopher Woods for helping us to set everything up and to Richard Pitts from Oracle. The authors also gratefully acknowledge the support of the EPSRC Digital Core Capability funding (EP/X034828/1) for the University of Bristol Digital Labs platform which provided high performance computing resources, as well as the support of all the staff involved. We would also like to thank the members of the Covert lab at Stanford University who have answered all our questions regarding the installation and use of the *E. coli* WCM, especially Dr. Travis Ahn-Horst and Dr. Gwanggyu Sun.

## Author contributions

I. M. G. developed the research concept (with assistance from L. M., W. P., Z. S. A., C. S. G. and T. E. G), built and trained the ML models, adapted the genome reduction algorithm, conducted the numerical simulations, analyzed the results, provided interpretation of the biological relevance and wrote the manuscript (with assistance from J. R-G., L. M., C. S. G. and T. E. G). K. S. compared the simulation results to the literature data on single gene essentiality and wrote most of the corresponding paragraphs of the manuscript. All authors reviewed the manuscript and approved the final version.

## Data and materials availability

The data used in this paper is stored using the storage system of the University of Bristol at https://data.bris.ac.uk/data/dataset/2wnzwzenfn4tt2iva9xxmb7ayh. The code used to generate the results presented in this study can be found at https://zenodo.org/records/13692990.

## Supplementary materials

Materials and Methods

Supplementary Text

Figs. S1 to S11

Tables S1 to S6

References *(43-68)*

Supplementary Table SE1-SE13

