## Supplementary text and methods for "Accelerated design of *Escherichia coli* reduced genomes using a whole-cell model and machine learning"

**This PDF file includes:**

Materials and Methods

Supplementary Text

Supplementary Figures S1 to S11

Supplementary Tables S1 to S6

**Other Supplementary Materials for this manuscript:**

Supplementary Tables SE1-SE13

### Materials and Methods

#### The *E. coli* whole-cell model simulations

In this study, we are using the single-cell version of the *E. coli* whole-cell model,[3] which can be thought of as a large system of Ordinary Differential Equations (ODEs) where the state variables of the ODEs correspond to different cellular states, and the differential equations correspond to cellular processes.[3] Simulating a cell is similar to numerical integration of the ODE system. The resources stored in cellular states must be shared with all cellular processes to ensure mass conservation.[3] In addition, the model has two main stochastic components: initializing RNA/protein synthesis and degrading RNA/protein. This is implemented by using a multinomial and Poisson distribution, respectively. The model is implemented in Python, using Cython and is available at `https://github.com/CovertLab/wcEcoli`. A version of the model available in March 2022 was used to run the simulations from in this study that produced the reduced genome EMine-737.

The model simulates the log phase growth of each cell and it is possible to do so for several generations. In this study, we used the default M9 minimal media supplemented with 0.4% glucose. For each cell simulation, the model outputs 219,207 time series after approximately 15 minutes of running time on a CPU. Given the large number of experiments needed for this study, the simulations were run in parallel on the University of Bristol's supercomputers BlueCrystal phase 4 and BluePebble, on a cluster in the cloud platform called Digital Labs, and on Oracle Cloud. The specifications of each of these systems can be found in Supplementary Table 12. Furthermore, the *E. coli* WCM uses Fireworks, a workflow management tool, to automate the execution of different jobs.

For the scope of this project, simulations were run for six generations, unless during this process a cell was not dividing in which case the simulations were sometimes stopped after four generations. Unless otherwise specified in the results section or in the supplementary material, the simulations were run for one repetition.

#### ML surrogate

A machine learning surrogate model is generally used to emulate the behaviour of a mechanistic model, with the added advantage that once trained, it can be used to make predictions several orders of magnitude faster than the original mechanistic model. The process for training such a machine learning model involves running the mechanistic model several times in order to generate the training and test sets. After that, the ML model is trained using this data, and once the accuracy is satisfactory, it can be used to replace the original mechanistic model in future simulations.[15] Some common techniques used to create surrogate models include regression models,[20,56] Gaussian processes,[22] support vector machines,[56] decision tree-based models[17,18] and neural networks.[16,18]

Typically, a machine learning surrogate model is trained on the parameters and/or initial conditions of the mechanistic model. However, due to the high number of parameters and variables of the WCM, this was not possible in this study due to the curse of dimensionality. Therefore, the ML surrogate is trained on the final state (just before division) of the first cell simulated for each repetition, and it predicts whether all the cells in the future five generations will divide. The advantage of this approach is that once it is trained, the mechanistic WCM has to be run for one generation, rather than six, and the data from this first generation is used to predict the fitness of all the cells in the future five generations.

In this paper we assess the accuracy of the ML surrogate using the F1-score metric, which combines the precision and recall scores of the models. We chose this metric due to its benefits when dealing with unbalanced data. A higher F1-score represents a better model.

#### Pearson correlation

Pearson correlation is an approach used to measure linear correlation between two variables $X$ and $Y$.[57] The Pearson correlation is calculated by

$$r = \frac{\sum(X_i - \overline{X})(Y_i - \overline{Y})}{\sqrt{\sum(X_i - \overline{X})^2 \sum(Y_i - \overline{Y})^2}}$$

where $X_i$ and $Y_i$ are the values of the two variables, $\overline{X}$ and $\overline{Y}$ are the means of the two variables and there are $n$ data points. A Pearson correlation coefficient of -1 indicates that the two variables are strongly negatively correlated, a Pearson correlation coefficient of 1 indicates that they are strongly positively correlated and a Pearson correlation coefficient equal to 0 indicates that there is no linear correlation.

#### Random forest

Random Forest is a popular supervised machine learning algorithm used for classification and regression problems.[58] It trains a large number of individual decision trees that operate as an ensemble. Each of these trees is trained on a random subset of the training examples and features. For classification tasks, the final predicted class is the one that was output by the majority of the decision trees. For regression models, each tree in the forest predicts a numerical value for a given observation based on the input features. The final output of random forest regressor is typically the average of all the tree predictions. This ensemble-based approach reduces overfitting and aids interpretability since it becomes possible to understand which features contributed to the prediction the most,[59] by averaging the gini impurity of the trees. Gini impurity in random forests measures the probability of misclassifying a randomly selected element if labeled according to the class distribution of a node. It serves as a splitting criterion in decision trees, prioritizing features that maximize Gini gain (the reduction in impurity achieved by splitting a node). For feature importance assessment in random forests, Gini importance averages the weighted impurity reduction across all trees for each feature.[38] Higher Gini importance scores indicate greater predictive power, as the feature effectively reduces class heterogeneity in splits.[38] This interpretability aspect is essential when used for biological applications since it helps users to understand why a system behaves in a specific way. Furthermore, compared to more complex deep learning models, random forest can be trained very quickly especially on small and medium data sets and it often has comparable accuracy. When there are many features available, but not many training examples, random forest is a more versatile and robust algorithm since its averaging process makes it less sensitive to noise in the data, reducing the likelihood of overfitting to noisy or irrelevant features and therefore providing an integrated feature selection methodology. The ML surrogate model based on random forest was built in Python using the scikit learn package.

#### XGBoost

XGBoost, or Extreme Gradient Boosting, is an advanced implementation of the gradient boosting algorithm designed for speed and performance.[60] It constructs an ensemble of decision trees where each tree corrects the errors of its predecessor, continuing this iterative process until a desired accuracy is achieved. XGBoost is known for handling missing values, incorporating regularization to prevent overfitting, and supporting parallel processing, which enhances its efficiency on large datasets. While XGBoost uses sequential tree building with regularization, random forest reduces overfitting by averaging predictions without built-in regularization. Random Forest can be advantageous due to its simplicity, requiring fewer hyperparameters and being less sensitive to settings, making it robust for quick models. It also offers better interpretability and naturally reduces variance, which can be beneficial in cases where overfitting is a concern.[60] The XGBoost model presented in the Supplementary Text of this manuscript was run in Python using the XGBoost open-source software library.

#### k-nearest neighbor

The k-nearest neighbors (KNN) algorithm is a non-parametric, supervised learning classifier that uses proximity to make predictions about the grouping of data points. It operates by storing the entire dataset and classifying new data points based on the majority class of their nearest neighbors, determined by a distance metric like Euclidean distance.[61] KNN does not involve a training phase; instead, it makes predictions directly from the dataset, which can be computationally expensive and memory-intensive. This simplicity and reliance on distance metrics make KNN sensitive to the choice of 'k' and the presence of outliers, and it performs poorly with high-dimensional data due to the curse of dimensionality.[61] Random forest and XGBoost are models that handle data more efficiently and make predictions faster. They are also more interpretable, less sensitive to noise and less likely to overfit the data.

#### Computational demand

Simulating the growth of one cell using the *E. coli* WCM takes approximately 15 minutes. Simulating cellular growth and division for 6 generations using the *E. coli* WCM involves simulating 63 cells (1 cell in generation 1, 2 cells in generation 2, 4 cells in generation 3 etc.). On a computer node this would take 945 minutes. Alternatively, using the ML surrogate proposed here, we remove the need to simulate 6 generation and instead we simulate only one generation of the WCM and use the ML surrogate to predict whether all cells in generations 2-6 will divide. This requires 15 minutes (the computational time required for one generation simulation since the ML algorithm runs in under 1 second), corresponding to a 98% reduction in computational time.

#### Minesweeper

Minesweeper is a four-stage algorithm inspired by laboratory fragment-cassette-fraction genome engineering,[29] which is similar to the divide-and-conquer paradigm. It was originally published in[11] to predict minimal genomes using the *M. genitalium* WCM, and it is freely available on `https://github.com/GriersonMarucciLab/Minesweeper`. A flow chart showing how Minesweeper works can be found in Figure S6 and the revised code is available at https://zenodo.org/records/13692990.

#### The new ML surrogate(that includes multiple gene knock outs in the training set)

The initial ML surrogate that was trained on single gene knock outs data was not performing very well on multiple gene knock outs and therefore a new ML model had to be trained using multiple gene knock outs. To achieve this we ran 6 generations simulations (using the WCM) where different permutations of the 5% segments run as part of stage 2 of Minesweeper are deleted. Figure S7A shows the segments that when deleted, the cell continues to divide for 6 generations. To generate the data to train the surrogate on multiple gene knock outs we split these segments such that permutations of the first six of them are used to test the new model and permutations of the last eight are used to train it (Figure S7B). Therefore, to the 738 single gene knock outs used to train the initial ML surrogate, we add 248 multiple gene knock out simulations (corresponding to a subset of all the permutations of the eight red segments from Figure S7B). Then we train a new random forest model using this data and we test its performance on the single gene knock outs test set, the double gene knock outs test set and the remaining multiple gene knock outs test set (corresponding to the 63 permutations of the blue segments from Figure S7B). The results are shown in Figure 3 and Table S4. This model corresponds to ML surrogate v2 in Figure 4.

A new ML model is trained one more time using another 59 knock outs from the fourth stage of Minesweeper (ML surrogate v3 in Figure 4). These correspond to multiple gene knock outs that were misclassified using the previous version of the surrogate. This new training stage was using data from the first two loops of stage four of Minesweeper and the model was used to make predictions for the last loop that outputted EMine-737. The F1-score of this model did not change when tested against the single, double and multiple gene knock out data sets.

### Gene ontology analysis

Gene Ontology (GO) is a major bioinformatics initiative that aims to unify the representation of gene and gene product attributes across all species. It provides an ontology of defined terms that represent gene product properties. The ontology covers three main domains: the Cellular Component domain, the Molecular function domain and the Biological Process domain. In this study, we used the databases and annotations from the Gene Ontology Consortium to perform our analysis.[62,63] The multiple false discovery rate used in this study is 0.05. The.obo file used to generate these results corresponds to the version available on 11th of June 2023.

In gene ontology (GO) analysis presented in this manuscript, the p-value represents a statistical measure used to determine whether a particular GO term is significantly enriched in a given set of genes compared to a control set (in this case the entire genome). Therefore, the p-value in the GO analysis represents the probability that the observed enrichment of a GO term occurred by random chance. A lower p-value suggests that the GO term is more significantly enriched in the gene set.

In this study, the GO terms were clustered based on the genes that they contain. Specifically, we used the Jaccard index to measure the similarity between sets of genes representing different ontology terms. Then we defined a distance matrix by using the formula: distance = 1 - jaccard_similarity. This transforms similarity values (where 1 = identical) to distance values (where 0 = identical). On this distance matrix we applied hierarchical clustering using a threshold of 0.9 to group terms containing common genes together. A threshold of 0.9 on the distance (which corresponds to a Jaccard similarity of 0.1) means we were clustering GO terms that have at least 10% gene overlap.

During our analysis, we noticed that genes included in some ontology categories differ slightly between EcoCyc[46] and the Gene Ontology Consortium database. One such example is the GO:0015871, which on EcoCyc includes the *betT* gene and in the database we use in this study it does not. We believe this emphasizes the need for more curated data sources in synthetic biology.

### Linear regression

Linear regression is an algorithm that models the relationship between a dependent variable and several independent variables. It is defined by a formula of the form $y = \beta_0 + \beta_1 x_1 + \beta_2 x_2 + ... + \beta_i x_i$, where $y_i$ is the dependent variable, $x_1, x_2, ..., x_n$ are the independent variables and $\beta_0 \beta_1, \beta_2 ...$ are the parameters (coefficients) to be estimated. The LinearRegression scikit-learn package in Python is estimating the parameters using the method of least squares, which aims to find the hyperplane that minimizes the sum of the squared residuals (the difference between the predicted and observed values).[64]

#### Experimental validation of single-gene knock out simulations

We compiled *in vivo* data from a publicly available CRISPR interference-based high-throughput screen that assessed ~3,400 nearly ubiquitous genes under three growth conditions in 18 representative *E. coli* strains,[40] we then focused on those consistently deemed essential in M9 minimal medium across all tested strains. This conservative approach ensured that only 'universally essential' genes were used to validate our predictions, accounting for the hybrid nature of the WCM genome, which integrates parameters defined across three different laboratory strains (K-12 MG1655, B/r, and BW25113).

Table SE12 shows the essentiality label for every gene in the experimental dataset and in the *in silico* dataset generated using the WCM and the ML surrogate.

### Supplementary Text

#### Feature engineering

To determine the number of cell divisions/generations to simulate *in silico*, we started by running the 1219 single gene knock out simulations for four generations. The results show that for 1101 of these genomes, all cells in four generations divided (non-essential knock out), and for 118 of them, at least one cell failed to divide (essential knock out). Among the 1101 cells, there are 43 that show growth defects, especially in their protein mass (Figure S1), suggesting that four generations of simulations might be too short to observe the phenotype of cells failing to divide. These 43 knock outs were simulated again for eight generations, together with another 61 randomly selected knock outs. The results confirmed that when simulating for eight generations, the 43 knock outs were essential.

The visual inspection of the masses of generation one cells corresponding to the 1101 non-essential gene knock outs shows that the protein mass of some cells stops growing. To identify these analytically we applied Euclidean k-means clustering on the time series. The results represented in Figure S1 show that the clustering algorithm is able to group all 43 cells in cluster 2.

We also inspected one randomly chosen gene knock out from cluster 2 (the deletion of alaS gene) and compared it to a gene knock out from cluster 1 (the deletion of the dfp gene). Figure S2 shows that when the *alaS* gene is knocked out the cells take longer to divide, probably due to a defect in growth. However, all cells in 4 generations divide, which is the reason why it was decided to run longer simulations of up to eight generations.

The inspection of the dynamics of the eight generations simulations shows that for 96% of the genomes, cells start failing to divide during the first six generations (Figure S3). Therefore, for training the ML surrogate cells should be allowed to divide for at least six generations.

The *E. coli* WCM is based on approximately 10,000 equations and 19,000 parameters.[3] The simulations output 219,207 time series for each cell. The size of the model does not allow for an ML surrogate to be trained on all of the WCM's initial conditions and/or parameters because of the curse of dimensionality[65] (i.e. there are more parameters/variables than training examples that are feasible to be generated using the WCM). To address this issue we decided to look at variables of the WCM that represent a summary of the state of the cell, such as the value of different macromolecular masses (e.g. the protein mass, RNA mass, DNA mass, membrane mass). In total, the model outputs 22 time series representing various masses, the growth rate and the cell's volume. We used the values of these variables at the beginning of the simulations (at $t$=0, generation 1) to train an ML model that predicts whether all cells in 6 generations will divide. The best F1-score obtained after training different ML algorithms was 0.76 using random forest (Table S1). This accuracy can be improved by 0.17 using the last time point of the macromolecular masses of the cell from the first-generation simulations (i.e. just before the cell divides) to predict the division of the cells from the first six generations (Table S1).

Pearson correlation is used to identify pairs of features with high correlation (i.e. with an absolute Pearson correlation coefficient above 0.95, an arbitrary chosen threshold, often used in the field, to represent strong linear correlation), and then to remove one of the redundant features, which helps to simplify the model without losing much information.[66] Furthermore, removing correlated features can improve interpretability, especially in linear models where multicollinearity affects coefficient estimation.[67] Based on Figure S4 we can see that the cell mass is perfectly correlated with the dry mass, the water mass, the cytosol mass and the cell volume. The cell mass is also highly correlated with the periplasm mass (the coefficient is > 0.95). This suggests that these features can be estimated based on the value of the other and therefore keeping one of them for further analysis is enough. Therefore, cell mass is kept as feature while dry mass, the water mass, the cytosol mass, the cell volume and the periplasm mass are removed and will not be considered in the correlation analysis for other features. Similarly based on the Pearson correlation coefficients in Figure S4, we see that the tRNA mass is highly correlated with the RNA mass and rRNA mass. Therefore, only the tRNA mass is kept as feature. The extracellular mass is highly correlated with the outer membrane mass, therefore only the extracellular mass is kept. The protein mass is highly correlated with the flagellum mass and the inner membrane mass, suggesting that all but the protein mass can be removed. This approach was used to reduce the number of features from 22 to 12.

#### Initial ML surrogate explainability and misclassifications

We further investigated the initial ML surrogate trained on single gene knock outs. First, we look deeper into the features used by the random forest classifier using their SHAP (SHapley Additive exPlanations) values. These are based on Shapley values from cooperative game theory.[68] In game theory, Shapley values help determine how much each player in a collaborative game contributes to the final outcome. In machine learning, SHAP values are used to explain the output of any machine learning model by assigning an importance value to each feature in a model. Features with positive SHAP values positively impact the model's output, while features with negative SHAP values negatively impact the model's output.

Figure S5 shows that the ML surrogate learns that a high cell mass has a negative impact on the model's output, meaning that the model is likely to predict that the knock out is non-deadly (the gene is non-essential), whereas for a lower cell mass the gene knocked out is likely to be essential. A similar, but less clear separation is observed for the DNA mass and the extracellular mass.

We also looked into the single gene knock outs that the ML surrogate model was not able to classify correctly. We employed gene ontology analysis on the set of these genes. The results can be found in the attached Supplementary Table SE11 and they show that out of the 30 unique misclassified genes, none were representing a full gene ontology. However, there was one GO category for which the knock out of 3 out of the 5 genes considered for training the ML surrogate were misclassified. These correspond to *thyA, prs* and *tmk* which are included in the biological process term representing nucleotide biosynthetic process. All the single knock outs of these 3 genes were classified as non-deadly, when according to the WCM they were deadly. It is likely that the ML surrogate misclassified these genes because the patterns representing a decrease in cell and DNA mass appear later in the simulations and are not present at the end of generation one.

**ML surrogate to predict growth rate**

An ML surrogate model that predicts the average growth rate of cells was built using the same data as in the case of the surrogate predicting cell division. We used the values of the 12 macromolecular masses (cell mass, growth, DNA mass, tRNA mass, extracellular mass, protein mass, projection mass, pilus mass, mRNA mass, small molecule mass, instantaneous growth rate, membrane mass) at the last time step of the first generation simulation to predict the average growth rate of the cells in generation 4. Generation 4 was chosen because for some simulations (where at least one cell was not dividing before generation 4), the simulations were stopped after 4 generations in order to save time. The training and test set were also split in the same way as in the case of the surrogate predicting cell division. The training set contains 369 different single gene knock outs, each repeated twice and representing a combination of simulations where cells divide successfully and not. The test set contains other 93 single gene knock outs, each repeated twice with diverse phenotypes. Figure S11 shows the distribution of the average growth rates (the predictors) for the test and train sets. It is possible to see that the average growth rate in generation 4 gives rise to 3 clusters, one representing simulations for which the cells were not growing, one representing cells that were growing slowly (average growth rate above 0, but bellow 0.5) and another one representing cells that grow faster (average growth rate above 0.5).

Before training the ML model, we standardized the macromolecular masses using the Standard-Scaler package from sklearn in Python. We tested the performance of three regression algorithms: random forest, XGBoost and linear regression. We tried several different combinations of hyper-parameter values for all algorithms and we chose the hyperparameters that were able to maximize the R-squared for the test set and minimize the difference between the R-squared of the training and test sets (in order to minimize overfitting). Table S6 shows the performance of all three models according to different metrics. The mean squared error (MSE) is a measure of the average squared difference between the estimated values and the actual values. The formula for this estimate is

$$\text{MSE} = \frac{1}{n}\sum_{i=1}^{n}(\hat{y}_i - y_i)^2.$$

The mean absolute error (MAE) is a measure of the average magnitude of the absolute errors between the predicted values and the actual values and it is defined by the formula

$$\text{MAE} = \frac{1}{n}\sum_{i=1}^{n}|\hat{y}_i - y_i|.$$

R-squared ($R^2$) is a statistical measure that represents the proportion of the variance for a dependent variable that is explained by an independent variable or variables in a regression model. The formula used to calculate the $R^2$ is

$$R^2 = 1 - \frac{\sum_{i=1}^{n}(y_i - \hat{y}_i)^2}{\sum_{i=1}^{n}(y_i - \overline{y})^2}.$$

For all these measures, $\hat{y}_i$ represents the predicted values, $y_i$ represents the actual values, $\overline{y}$ represents the mean of the actual values and $n$ is the number of data points considered.[64]

According to the results in Table S6, the performance of the models tested is similar, with a MSE on the test set between 0.0158 (for the random forest) and 0.0178 (for linear regression), a MAE for the test set between 0.0606 (for random forest) and 0.0732 (for linear regression) and a coefficient of determination ($R^2$) for the test set between 0.8922 (for random forest) and 0.8784 (for linear regression). These results suggest that random forest performs the best, while linear regression performs the worst. However, all algorithms are able to successfully predict growth rate and can be used as ML surrogates.

### Supplementary Figures and Tables

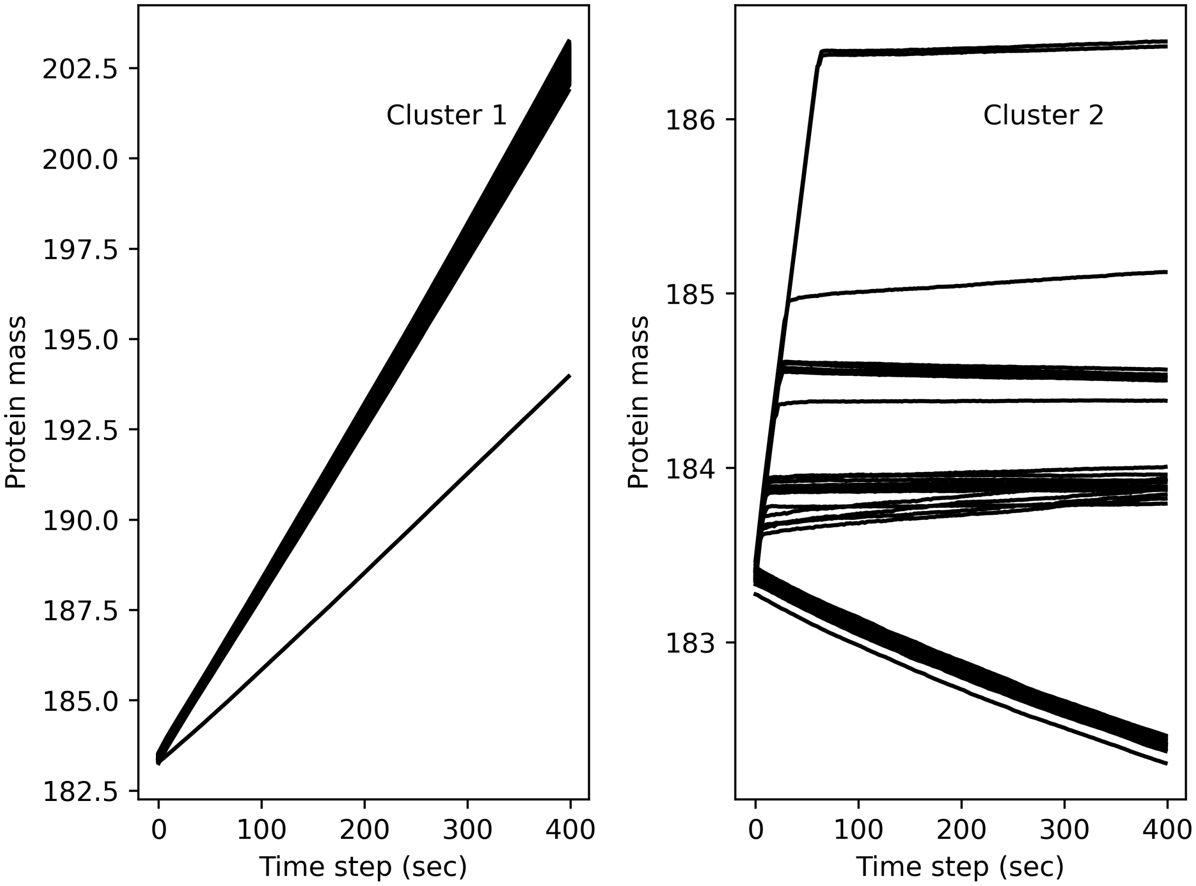


**Figure S1**: **First generation cells from non-essential (based on 4 generations simulations using the WCM) single gene knock out simulations.** Cluster 2 contains 43 cells for which the protein mass does not grow as expected. This suggests that the cells will probably stop dividing if their corresponding simulations are run for more than 6 generations. As shown in the main text, this is indeed the case.

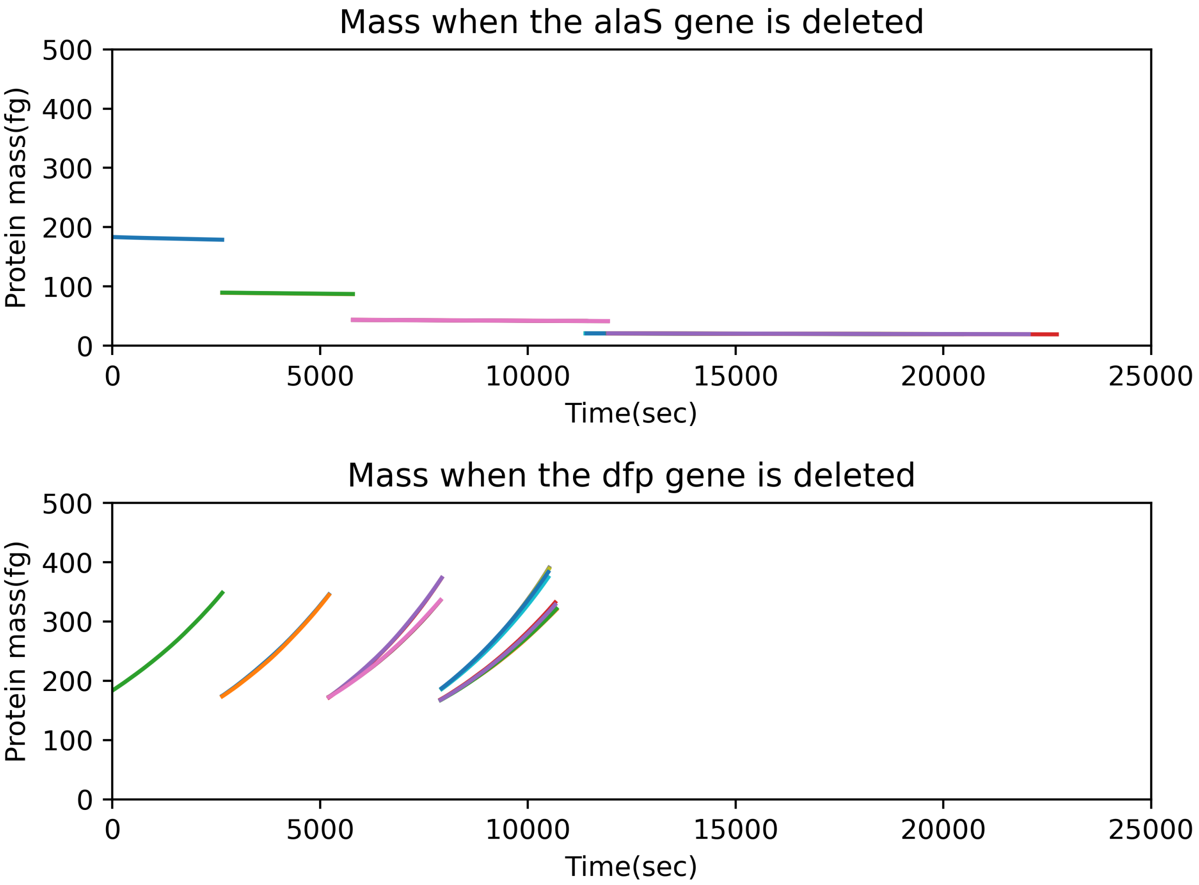


**Figure S2**: **The protein mass of cells grown for 4 generations upon single gene deletions.** The alaS gene deletion exhibits a defect in protein mass, while the dfp gene does not.

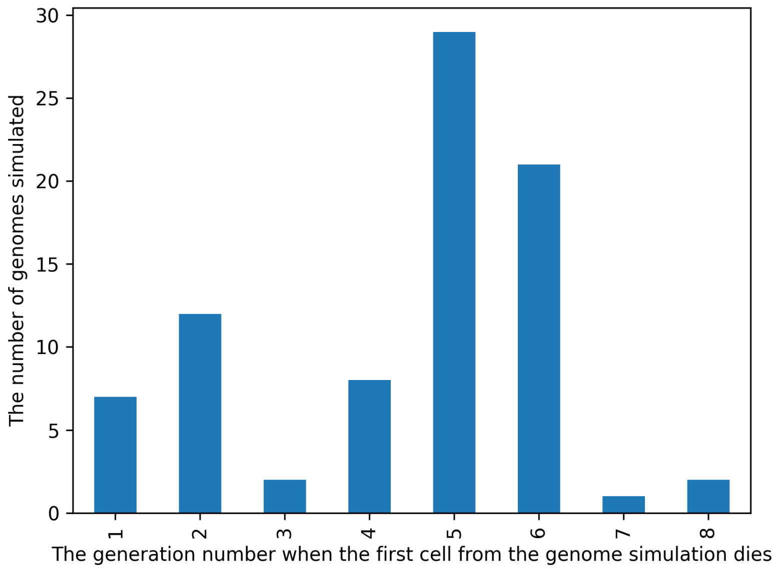


**Figure S3**: **A histogram showing the generation when the first cell in each genome simulated for 8 generations fails to divide.** For most knock outs the cells start failing to divide in generations 5 and 6. This suggests that we need to let the cells divide for at least 6 generations. This will cause some loss of information from the cells that fail to divide later, but given the high number of cells to simulate in generations 7 and 8 (64 and 128 respectively) and the high computational demand of the model, we will assume we can stop simulating after 6 generations.

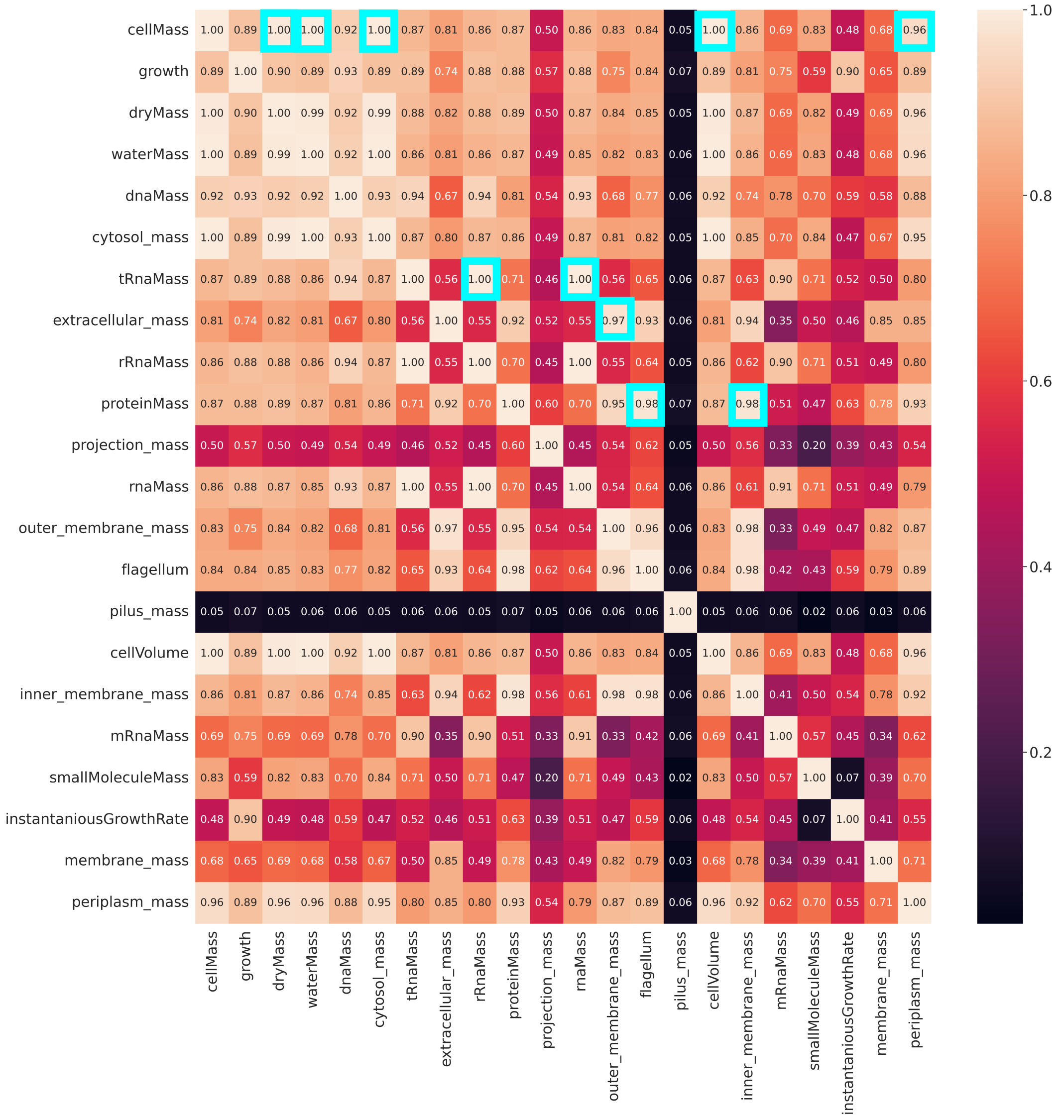


**Figure S4**: **Correlation matrix for the macromolecular masses at the last time step before division of generation 1 cells.** The numbers inside the matrix represent the absolute value of the Pearson correlation coefficient between each row-column pair. A Pearson correlation of 1 means that the two features are perfectly linearly correlated. The principal diagonal of the matrix only contains ones because all features are perfectly correlated to themselves. The blue squares highlight the columns (features) that were removed due to a correlation coefficient higher than 0.95. Once a column is highlighted for removal due to high correlation, it is not considered further for correlation calculation against other rows.

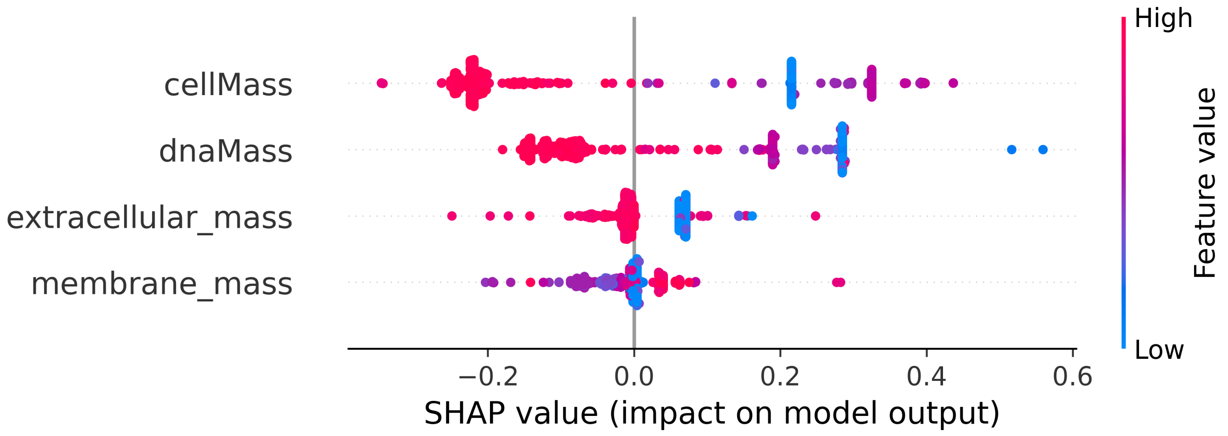


**Figure S5**: **SHAP values corresponding to the features of the initial ML surrogate.**

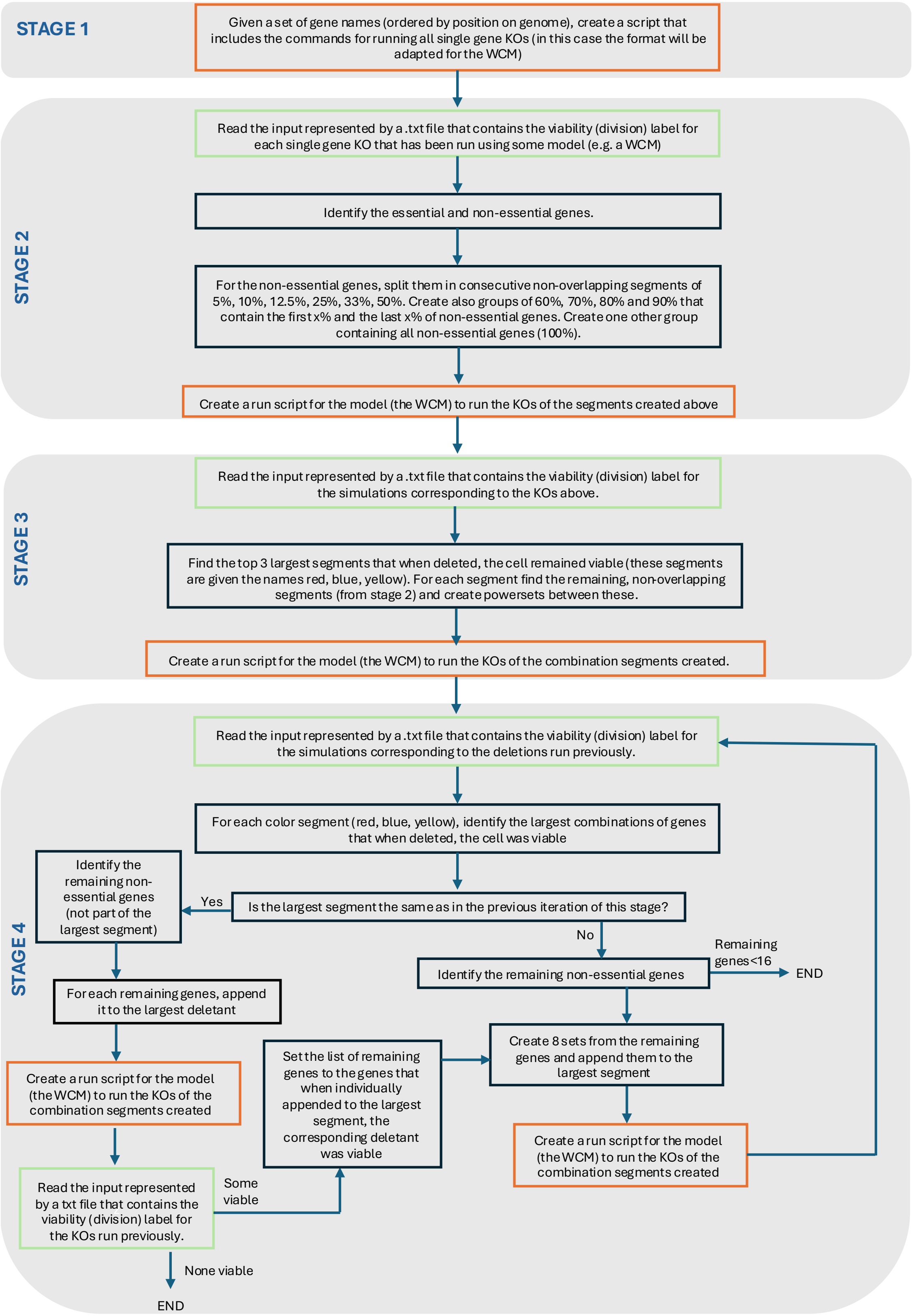


**Figure S6**: **Flowchart showing how Minesweeper works.** The logic behind the algorithm is also presented in Figure 4.

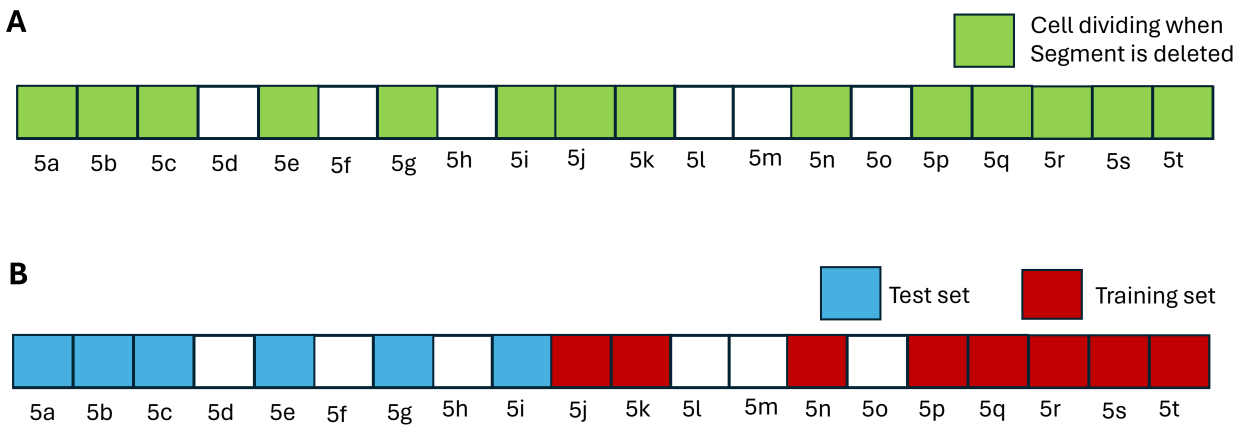


**Figure S7**: **Visual representation of the segments from stage 2 of Minesweeper.** (A) The segments of non-essential genes that when removed the cell divided for 6 generations. These deletions correspond to stage 2 of Minesweeper. (B) The split between these segments to obtain the training and test data for the ML surrogate trained on multiple gene knock outs as well.

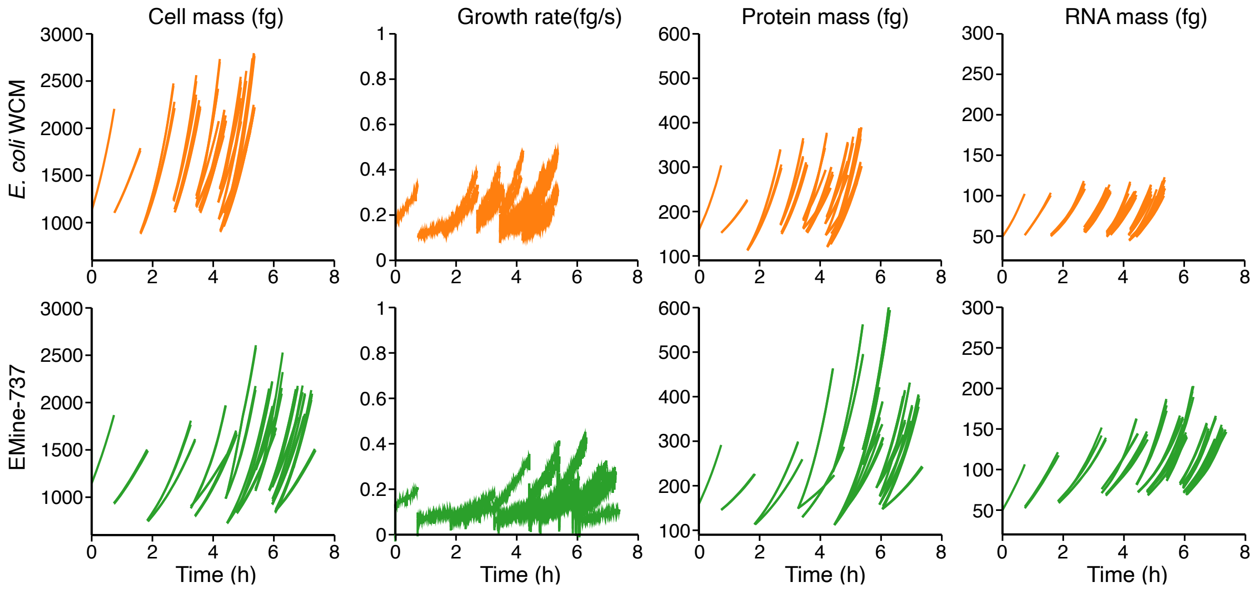


**Figure S8**: **An example of the WCM output for one repetition of an EMine-737 genome and one repetition of a wildtype genome.** The EMine-737 (first row) and wildtype (second row) genomes are allowed to divide for 6 generations, with each cell division producing two daughters. Each line in the graphs represents the cell mass, growth rate, protein mass or RNA mass of a cell.

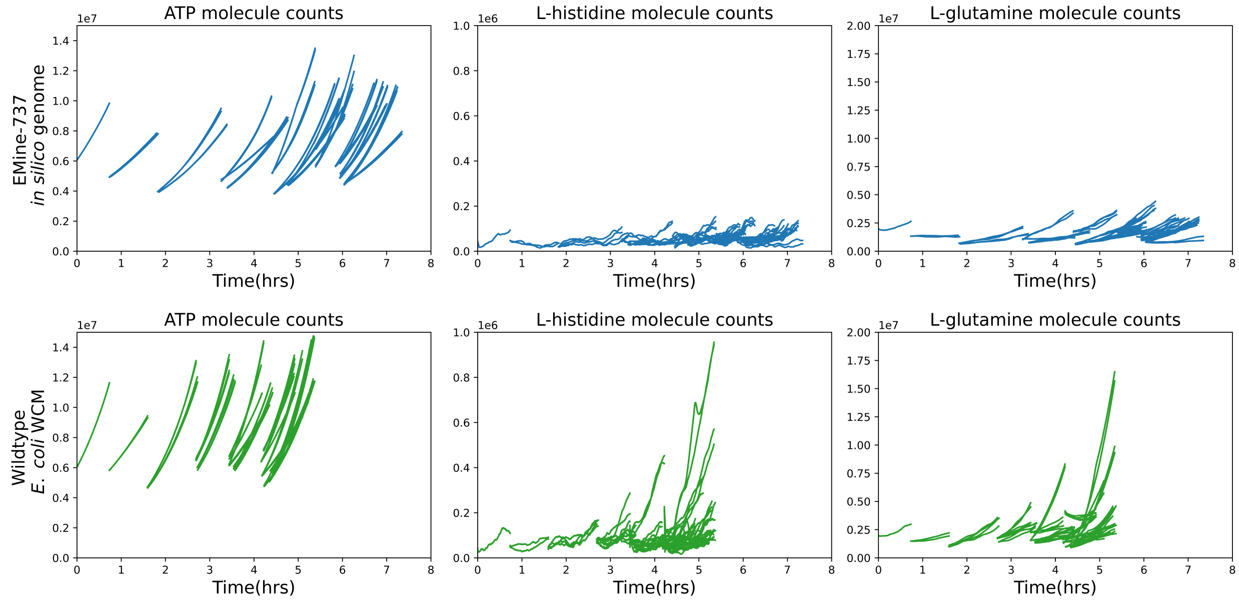


**Figure S9**: **A comparison between the molecule counts of ATP, L-histidine and L-glutamine between wildtype cells and EMine-737 cells**. The figure shows one example of an EMine-737 cell that divides 5 times and one example of a wildtype cell that divides 5 times. Each line in the graphs represents the corresponding molecule counts for one cell.

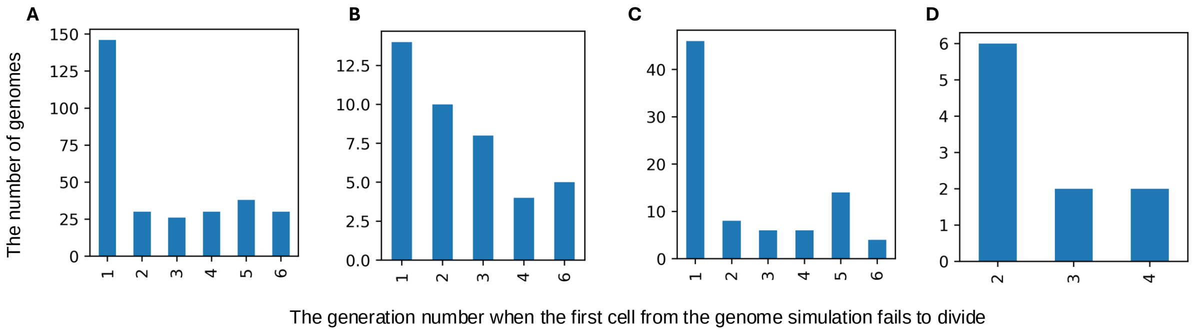


**Figure S10**: **Histogram showing the generation when the first cell in the simulation fails to divide.** Panel **A** represents the distribution of the generation when the first cell fails to divide for the data used to trained the initial ML surrogate. Panel **B** represents the distribution of the generation when the first cell fails to divide in the simulations from the training set that misclassify essential genes. The missing generations from the x axis suggest that there was no misclassified simulation when the first cell death occurred in that generation. Panel **C** represents the distribution of the generation when the first cell fails to divide for the data used to test the initial ML surrogate. Panel **D** represents the distribution of the generation when the first cell fails to divide in the simulations from the test set that misclassify essential genes. The missing generations from the x axis suggest that there was no misclassified simulation when the first cell death ocured in that generation.

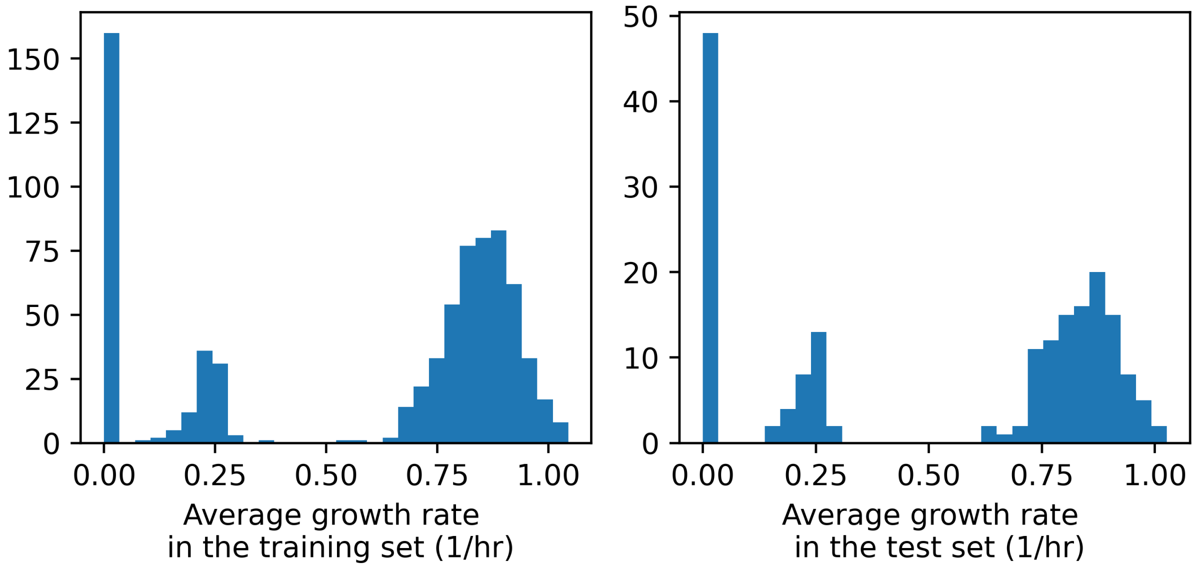


**Figure S11**: **The distribution of the average growth rate for the data used to train and test the initial ML surrogate.** The growth rates on the left come from generation 4 of the 738 (369*2 because each knock out simulations was repeated twice) gene knock out simulations used to train the ML surrogate for growth rate. The growth rates on the right come from generation 4 of the 186 (93*2 because each knock out simulations was repeated twice) gene knock out simulations used to train the ML surrogate for growth rate.

| Algorithm | F1-score training data | F1-score test data | Feature description |
|---|---|---|---|
| Random Forest | 0.75 | 0.76 | The value of the 22 macromolecular masses outputted by the WCM at the **first** time step of the first generation simulation. |
| XGBoost | 0.73 | 0.73 | |
| K-Nearest Neighbour | 0.71 | 0.7 | |
| Random Forest | 0.92 | 0.93 | The value of the 22 macromolecular masses outputted by the WCM at the **last** time step of the first generation simulation. |
| XGBoost | 0.91 | 0.92 | |
| K-Nearest Neighbour | 0.91 | 092 | |

**Table S1**: **The performance of the machine learning models used to identify the data that should be used to train the ML model.** 22 macromolecular masses (total cell mass, growth, dry mass, water mass, dna mass, cytosol mass, tRna mass, extracellular mass, rRna mass, protein mass, projection mass, rna mass, outer membrane mass, flagellum, pilus mass, cell volume, inner membrane mass, mRna mass, small molecule mass, instantanious growth rate, membrane mass, periplasm mass) are used as features. The top part of the table presents the F1-score of each algorithm when the first time step of the WCM simulations is used to train the machine learning model and the lower part of the table represents the same metrics when the last time step of generation 1 simulations is used as features.

| Data set | Number of generations | Number of seeds/ repetitions | Number of unique KOs in the training set | Number of unique KOs in the test set |
|---|---|---|---|---|
| Single gene knock outs | 6 or 4 | 2 | 369 | 93 |
| Multiple gene knock outs | 6 | 1 | 307 (for the model trained during the last stage of Minesweeper) | 63 |
| Double gene knock outs | 6 | 1 | 0 | 47 |

**Table S2**: **Simulated data used to train and test the ML surrogates.** In the case of the single gene knock out, most simulations were run for minimum 6 generations, except for the essential gene knock outs identified based on the four generations simulations described at the beginning of this section.

| Data set | F1-score | Precision | Recall | Accuracy |
|---|---|---|---|---|
| Training on single KOs | 0.92 | 0.86 | 0.99 | 0.94 |
| Test single KOs | 0.93 | 0.88 | 0.97 | 0.94 |
| Test double KOs | 0.93 | 0.88 | 1 | 0.90 |
| Test multiple gene KOs | 0.82 | 0.86 | 0.79 | 0.78 |

**Table S3**: **The performance metrics for the initial ML surrogate trained only on single gene knock outs.** The confusion matrices corresponding to these metrics can be found in Figure 2A, the features used by this model are shown in Figure 2B.

| Data set | F1-score | Precision | Recall | Accuracy |
|---|---|---|---|---|
| Training on single and multiple KOs | 0.93 | 0.88 | 0.99 | 0.94 |
| Test single KOs | 0.93 | 0.88 | 0.97 | 0.94 |
| Test double KOs | 0.93 | 0.86 | 1 | 0.89 |
| Test multiple gene KOs | 0.88 | 0.78 | 1 | 0.87 |

**Table S4**: **The performance metrics for the new ML surrogate trained on single and multiple gene knock outs.** The confusion matrices corresponding to these metrics can be found in Figure 3A, the features used by the model are shown in Figure 3B.

| Accuracy | Precision | Recall | F1 Score |
|---|---|---|---|
| 0.871 | 0.904 | 0.943 | 0.923 |

**Table S5**: **Metrics assessing the agreement between experimental *in vivo* measured gene essentiality and *in silico* measured gene essentiality outputted by the ML surrogate.**

| Algorithm name | MSE train | MSE test | MAE train | MAE test | $R^2$ train | $R^2$ test |
|---|---|---|---|---|---|---|
| Random forest | 0.0103 | 0.0158 | 0.0593 | 0.0606 | 0.9269 | 0.8922 |
| XGBoost | 0.0102 | 0.0162 | 0.0684 | 0.0711 | 0.9273 | 0.8899 |
| Linear Regression | 0.0124 | 0.0178 | 0.0712 | 0.0732 | 0.9113 | 0.8784 |

**Table S6**: **Performance of the ML surrogate trained to predict growth rate.** For each of the five algorithms we tested, we measure the Mean Squared Error (MSE), the Mean Absolute Error (MAE) and the coefficient of determination ($R^2$) for the training and test set.

**Caption for Supplementary Table SE1. The list of genes that were knocked out to obtain the data that was used to train and test the initial ML surrogate.**

**Caption for Supplementary Table SE2. The data used to train the new ML surrogate representing multiple gene knock out simulations.** The table represents the value of each macromolecular mass at the end of generation 1, the corresponding label (deadly_gene/non_deadly), the path to the cell's simulation data, the ID of the segments (from Minesweeper stage 2) used to generate the KOs and the genes that were knocked-out.

**Caption for Supplementary Table SE3. A list of all the genes modeled in the WCM and their essentiality label according to the ML surrogate.**

**Caption for Supplementary Table SE4. A list of the deletion segments (and the corresponding genes) proposed by Minesweeper in stage 2 that when removed all cells in 6 generations divided.**

**Caption for Supplementary Table SE5. A list of the genes deleted in EMine-737 and a list of the remaining non-essential genes (according to the surrogate and the WCM).**

**Caption for Supplementary Table SE6. The genes used to perform the comparison between EMine-737 and the MS56 genome from the literature.** The comparison workbook contains all genes in MG1655 and whether they were removed in EMine-737 or MS56, as well as whether they are modelled by the WCM. The EMine-737 deleted non-metabolic workbook contains the non-metabolic genes deleted from EMine-737.

**Caption for Supplementary Table SE7. All the GO terms that contained genes included in EMine-737 and in MS56 and the corresponding genes.** The table contains columns corresponding to the GO term ID (GO), a short description of the term (term), the class that the term belongs to (class), the p-value (p), the number of genes from our study that correspond to that GO term (n_genes), the total number of genes in our study included in the analysis (n_study), the number of genes in that GO term (n_go), the genes from each GO term that are included in our anlaysis (study_genes) and the proportion of genes from the GO term that are also present in our analysis n_genes/n_go (per).

**Caption for Supplementary Table SE8. All the GO terms that contained genes included in EMine-737, but deleted from MS56 and the corresponding genes.** The table contains columns corresponding to the GO term ID (GO), a short description of the term (term), the class that the term belongs to (class), the p-value (p), the number of genes from our study that correspond to that GO term (n_genes), the total number of genes in our study included in the analysis (n_study), the number of genes in that GO term (n_go), the genes from each GO term that are included in our anlaysis (study_genes) and the proportion of genes from the GO term that are also present in our analysis n_genes/n_go (per).

**Caption for Supplementary Table SE9. All the GO terms that contained genes removed from EMine-737 and from MS56 and the corresponding genes.** The table contains columns corresponding to the GO term ID (GO), a short description of the term (term), the class that the term belongs to (class), the p-value (p), the number of genes from our study that correspond to that GO term (n_genes), the total number of genes in our study included in the analysis (n_study), the number of genes in that GO term (n_go), the genes from each GO term that are included in our anlaysis (study_genes) and the proportion of genes from the GO term that are also present in our analysis n_genes/n_go (per).

**Caption for Supplementary Table SE10. All the GO terms that contained genes removed from EMine-737, but kept in MS56 and the corresponding genes.** The table contains columns corresponding to the GO term ID (GO), a short description of the term (term), the class that the term belongs to (class), the p-value (p), the number of genes from our study that correspond to that GO term (n_genes), the total number of genes in our study included in the analysis (n_study), the number of genes in that GO term (n_go), the genes from each GO term that are included in our anlaysis (study_genes) and the proportion of genes from the GO term that are also present in our analysis n_genes/n_go (per).

**Caption for Supplementary Table SE11. Information on the genes with misclassified viability by the initial ML surrogate when compared to the WCM.** The Misclassified genes workbook contains the genes that were misclassified by the initial ML surrogate. The 'GO analysis of miscl. genes' workbook contains the GO analysis corresponding to the misclassified genes.

**Caption for Supplementary Table SE12. The labels used to compare the experimental essentiality data to the ones produced using the WCM and ML surrogate.**

**Caption for Supplementary Table SE13. Information regarding the computational infrastructure used to run the simulations.**
